# Towards a catalog of pome tree architecture genes: the draft ‘d’Anjou’ genome (*Pyrus communis* L.)

**DOI:** 10.1101/2021.11.17.467977

**Authors:** Huiting Zhang, Eric K. Wafula, Jon Eilers, Alex E. Harkess, Paula E. Ralph, Prakash Raj Timilsena, Claude W. dePamphilis, Jessica M. Waite, Loren A. Honaas

## Abstract

The rapid development of sequencing technologies has led to a deeper understanding of horticultural plant genomes. However, experimental evidence connecting genes to important agronomic traits is still lacking in most non-model organisms. For instance, the genetic mechanisms underlying plant architecture are poorly understood in pome fruit trees, creating a major hurdle in developing new cultivars with desirable architecture, such as dwarfing rootstocks in European pear (*Pyrus communis*). Further, the quality and content of genomes vary widely. Therefore, it can be challenging to curate a list of genes with high-confidence gene models across reference genomes. This is often an important first step towards identifying key genetic factors for important traits. Here we present a draft genome of *P. communis* ‘d’Anjou’ and an improved assembly of the latest *P. communis* ‘Bartlett’ genome. To study gene families involved in tree architecture in European pear and other rosaceous species, we developed a workflow using a collection of bioinformatic tools towards curation of gene families of interest across genomes. This lays the groundwork for future functional studies in pear tree architecture. Importantly, our workflow can be easily adopted for other plant genomes and gene families of interest.

## Introduction

Advancements in plant genome sequencing and assembly have vigorously promoted research in non-model organisms. In horticultural species, new genome sequences are being released every month ^1–6^. These genomes have broadened our understanding of targeted cultivars and provided fundamental genomic resources for molecular breeding and more in-depth studies of economically important crop traits such as those involved in plant architecture. Although many gene families have been identified as important for architectural traits, such as dwarfing, weeping, and columnar growth^7^, the study of these genes and their functionality in new species is still hampered by inaccurate information about their gene models or domain structures, and the frequent lack of 1:1 orthology between related genes of different study species. Sequencing and annotating a diversity of related genomes are crucial steps for obtaining this level of information.

Crops, most of which have gone through more than ten thousand years of domestication to meet human requirements, have a wide diversity in forms, sometimes even within the same species^8^. One such example is in the *Brassica* species, where *B. rapa* encompasses morphologically diverse vegetables such as Chinese cabbage, turnips, and mizuna; and cabbage, stem kale, and Brussels sprouts are the same biological species, *B. oleracea*. Therefore, a single reference genome does not represent the complex genome landscape, or pan-genome, for a single crop species. To understand the genetic basis of the diverse *Brassica* morphotypes, many attempts have been made to explore the genomes of *Brassica*^8–12^. In one of those attempts, genomes from 199 *B. rapa* and 119 *B. oleracea* accessions were sequenced and analyzed using a comparative genomic framework^10, 12^. Genomic selection signals and candidate genes were identified for traits associated with leaf-heading and tuber-forming morphotypes. Compared to *Brassica*, pome fruits may not appear to have as much diversity in their vegetative appearance, but they do have great diversity in terms of fruit quality, rootstock growth and performance, and post-harvest physiology. However, genome studies and pan-genome scale investigations in pome fruits are still in their infancy. In cultivated apple (*Malus domestica*), genomes of three different cultivars^13–16^ have been published, providing resources to study: 1) small (SNPs and small InDels) and large scale (chromosome rearrangements) differences that can help explain cultivar diversity, and 2) gene content differences that may contribute to cultivar specific traits. However, genomic resources for European pear (*Pyrus communis*) cultivars are limited to just two published genomes^17, 18^ from a single cultivar, ‘Bartlett’. More European pear genomes will afford new perspectives that help us understand shared and unique traits for important cultivars in *Pyrus*, as well as other Rosaceae.

Besides understanding large scale genomic characteristics, new genomes also provide rich resources for reverse genetic studies^19, 20^. To obtain the actual sequence of a target gene, reverse genetic approaches in the pre-genome era relied on sequence and domain homology and technologies such as RACE PCR^21^, which could be challenging and time consuming. Alternatively, in species with high-quality reference genomes, the annotation is generally considered to contain all the genes and target genes could ideally be identified with a sequence similarity search (*i.e.,* BLAST). However, reports of annotation errors, such as imperfect gene models and missing functional genes are very common^22–24^. Another complicating factor is that duplication events (*i.e.,* whole genome duplication, regional tandem duplication) and polyploidy occur in the majority of flowering plants, including most crop species, posing substantial challenges to genome assembly and annotation^25^. Moreover, instances of neofunctionalization and subfunctionalization occur frequently following duplication events^26^, which sometimes will result in large and complex gene families^27, 28^. Therefore, a one-to-one relationship between a gene in a model organism and its ortholog in other plant species, or even between closely related species and varieties, is rare^29^. Without understanding the orthology and paralogy between members of a given gene family, it is difficult to translate knowledge of a gene in a model organism to another species of interest.

In the present study, we assembled a draft genome for the European pear cultivar ‘d’Anjou’, improved the current ‘Bartlett’ assembly (*i.e.,* Bartlett.DH_V2), and developed a workflow that allows highly efficient target gene identification in any plant genome of interest. We used our workflow to curate and improve gene models for architecture-related genes from both the polished Bartlett.DH_v2 and the d’Anjou genomes. Importantly, we recovered many genes that were missing from gene families of interest (50 genes in the cultivar ‘Bartlett’) and corrected errors in others across the genus *Pyrus*. This work demonstrates that the integration of comparative genomics and phylogenomics can facilitate and enhance gene annotation, and thus gene discovery, in important plant reference genomes.

## Results

### The draft d’Anjou genome

#### Genome assembly

We generated approximately 134 million paired-end reads from Illumina HiSeq and a total of 1,054,992 PacBio continuous long reads (CLR) with a read length N50 of 20 Kb, providing an estimated 67-fold and 21-fold coverage respectively of the expected 600 Mb *Pyrus communis* genome^18^. Additionally, approximately 468 million 2 x 150 bp paired reads (∼234-fold coverage) with an estimated mean molecule length (linked-reads) of 20 kb were generated using 10x Genomics Chromium Technology (Supplementary Table 1). The final meta-assembly, generated with a combination of the three datasets, contains 5,800 scaffolds with a N50 of 358 Kb (Table 1). The cleaned contigs and scaffolds were ordered and oriented into 17 pseudochromosomes guided by the reference genome, *Pyrus communis* ‘Bartlett.DH_v2’^17^.

**Table 1.** Comparison of genome assembly and annotation, and orthogroups among *Pyrus communis* genotypes.

| Characteristics | Bartlett_v1 | Bartlett.DH_v2 | d'Anjou_v1 |
| --- | --- | --- | --- |
| Assembly |  |  |  |
| Assembly size (Mb) | 600 | 507.7 | 600 |
| Number of scaffolds | 142,083 | 592 | 5800 |
| Scaffold N50 | 88 Kb | 8.1 Mb | 358.88 Kb |
| Pseudochromosomes | 17 | 17 | 17 |
| Annotation |  |  |  |
| Predicted gene number | 43,419 | 37,445 | 45,981 |
| BUSCO | 96.5% | 96.5% | 96.6% |
| Mean CDS length | 1209 | 1120 | 1343 |
| Gene family classification |  |  |  |
| Percentage of genes classified into pear orthogroups | 76.2 | 76.2 | 80.4 |
| Percentage of pear orthogroups containing genes | 93.7 | 81 | 90.7 |
| Number of 26Gv2 orthogroups containing genes | 9878 | 9668 | 9837 |

Next, we compared the d’Anjou meta-assembly to two published reference assemblies of Bartlett^17, 18^ to assess assembly contiguity, completeness, and structural accuracy. The Benchmarking Universal Single-Copy Ortholog (BUSCO)^30^ analysis showed that the d’Anjou genome captured 96.6% complete genes in the Embryophyta gene sets, comparable to the reference genomes (Table 1, Supplementary Table 2). Furthermore, synteny comparisons between the draft d’Anjou genome and the reference Bartlett.DH_v2 genome showed high collinearities at both whole-genome and chromosomal levels (Fig. 1a and Supplementary Fig. 1).

**Fig 1.**
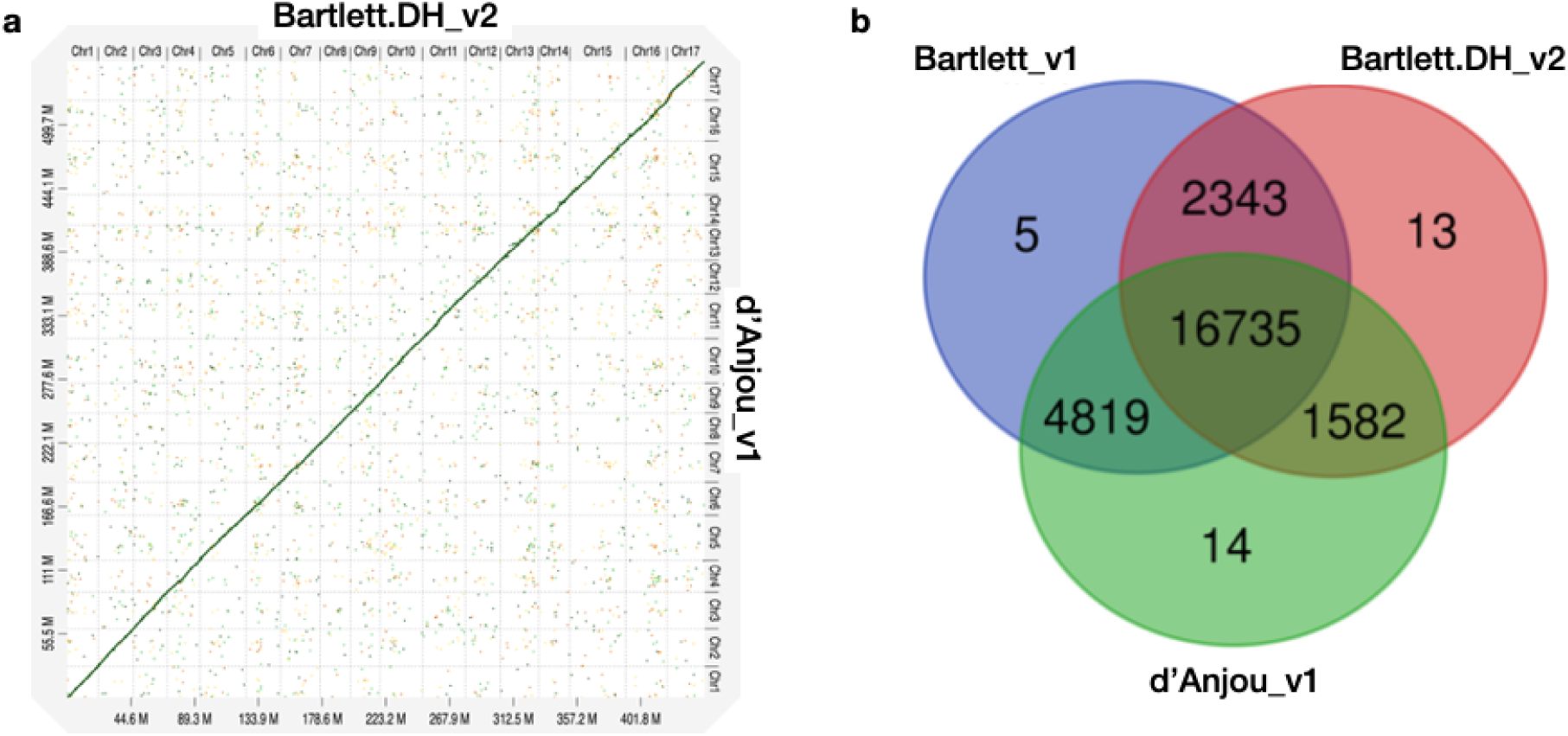
Characterization of the d’Anjou_v1 genome and protein orthology among European pears. a. Dot plot of genome alignment of Bartlett.DH_v2 (x axis) and d’Anjou_v1 (y axis). **b.** Overlap and distinctiveness of gene annotations among three *Pyrus communis* genotypes, Bartlett_v1, Bartlett.DH_v2, and d’Anjou.

#### Annotation

Combining information such as *de novo* transcriptome assembly, homologous proteins of closely related species, and protein-coding gene annotations from the two Bartlett genomes, we identified a total of 45,981 protein coding genes in d’Anjou (Table 1). Of those putative genes 76.63% were annotated with functional domains from Pfam^31^ and the remaining are supported by annotation evidence, primarily d’Anjou RNA-Seq reconstructed transcript^32^. These results indicate that we captured a large majority of the gene space in the d’Anjou genome. This affords a range of analyses including gene and gene family characterization, plus global-scale comparisons with other Rosaceae including the ‘Bartlett’ cultivar.

### Comparison among three European pear genomes

To study the shared and genotype-specific genes among the three European pear genomes, we constructed 25,511 protein clusters, comprising 77.71% of all the genes. While numbers of predicted genes from the Bartlett_v1 and d’Anjou genomes may be overestimated due to the presence of alternative haplotype segments in the assembly caused by high heterozygosity^17^, this should have very little effect on orthogroup circumscription. Further, the process of creating a double haploid reduces genome heterozygosity, but should retain estimates of orthogroup content. Hence, we formulated the following hypotheses: 1) a large majority of gene families are shared by all three genotypes; 2) few genotype-specific gene families are present in each genome; 3) the commercial ‘Bartlett’ genotype and the double haploid “Bartlett’ genotype (roughly version 1.0 and 2.0 of this genome, respectively) should have virtually identical gene family circumscription; and 4) we should detect very few gene families that are unique to either ‘Bartlett’ genome and shared with ‘d’Anjou’. The protein clustering analysis results (Table 1, Fig. 1b) support our hypotheses 1 and 2: 65.60% of the orthogroups contain genes from all three genotypes and only 0.12% of the orthogroups are species-specific. However, among the 8,744 orthogroups containing genes from two genotypes, more than half (55.11%) are shared between d’Anjou and Bartlett_v1, 18.10% are shared by d’Anjou_v1 and Bartlett.DH_v2, and only 26.80% are shared between the two Bartlett genomes, which does not support hypotheses 3 and 4.

To better understand why these hypotheses lacked support, we took a broader look at gene family content by comparing a collection of Rosaceae genomes, including the pear genomes in question. We assigned all the predicted protein coding genes from genomes of interest^13–15, 17, 18^ to orthogroups constructed with a 26-genome scaffold, covering most of the major lineages of land plants (supplementary Fig. 2). Out of the 18,110 orthogroups from this database, *Prunus persica*, a rosaceous species included in the genome scaffold, has representative genes in 10,290 orthogroups. Genes from most apple and pear genomes (Bartlett_v1, d’Anjou_v1, *Malus domestica* HFTH_v1.0, *M. domestica* GDDH13_v1.1, *M. domestica* Gala_v1.0, *M. sieversii*_v1.0, *M. sylvestris*_v1.0) are present in more than 9,800 orthogroups, however, genes from Bartlett.DH_v2 were only found in 9,688 orthogroups (Table 1 & Supplementary Table 3). These results suggest there are many genes not annotated in the Bartlett.DH_v2 genome.

### Genome-Wide identification of selected architecture genes

#### A selection of architecture genes

With this new comparative genomic information, our next steps were two-fold: first, to leverage information from the three European pear genomes and other available Rosaceae genomes, to identify and improve a set of tree architecture-related gene models of interest, and second, to use these architecture gene families as a test case to investigate potential issues in the Bartlett.DH_v2 genome. Many aspects of tree architecture are important for improving pear growth and maintenance, harvest, ripening, tree size and orchard modernization, disease resistance, and soil microbiome interaction. Traits of interest include dwarfing and dwarfism, root system architecture traits, and branching and branch growth. We selected key gene families known to be involved (Table 2)^33–64^, particularly those that have been previously shown to influence architectural traits in fruit trees. The identification of genes within these families, as well as their genomic locations, correct gene models, and domain conservation, is an important early step in testing and understanding their relationships and functions.

**Table 2.**
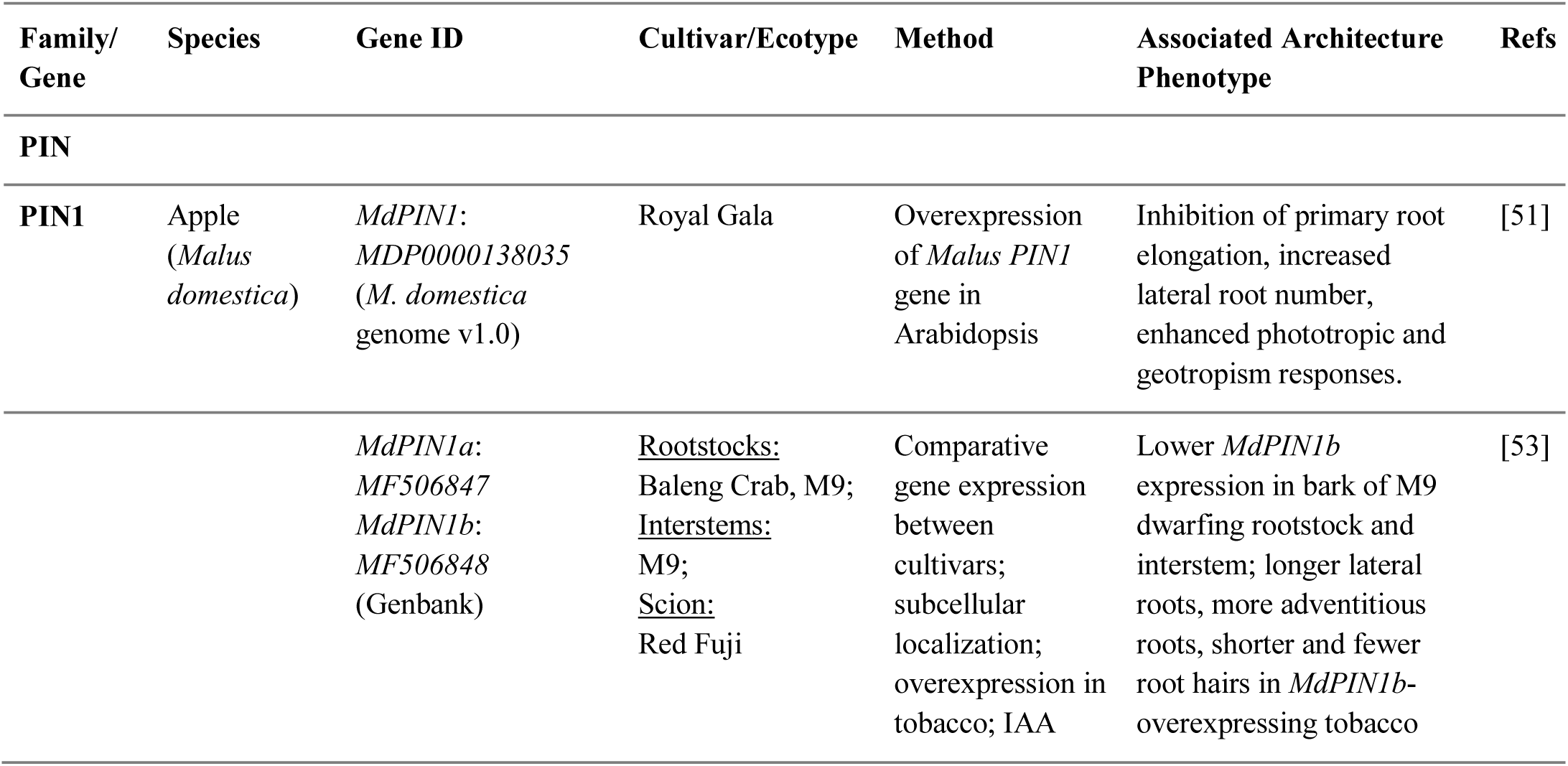
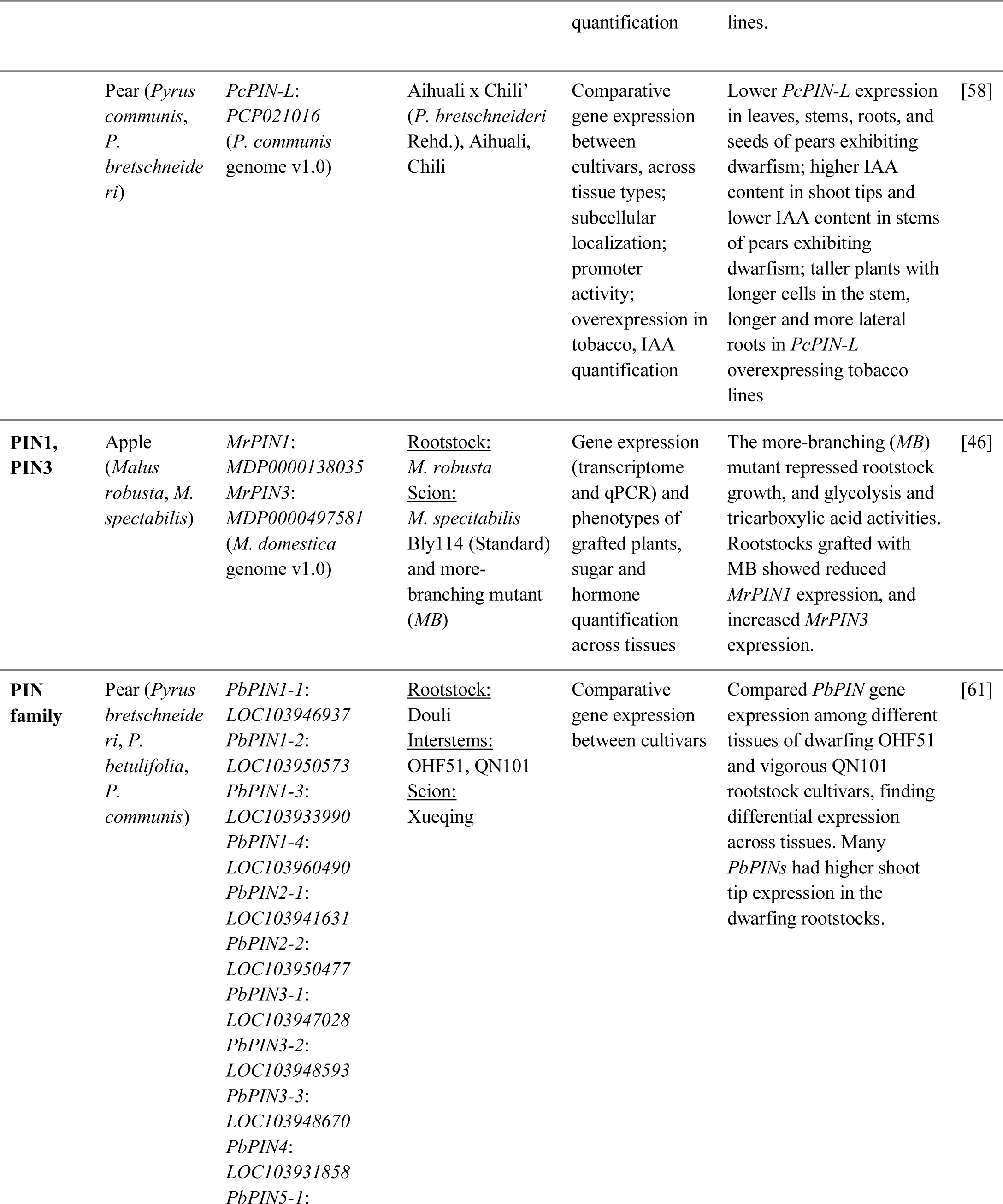
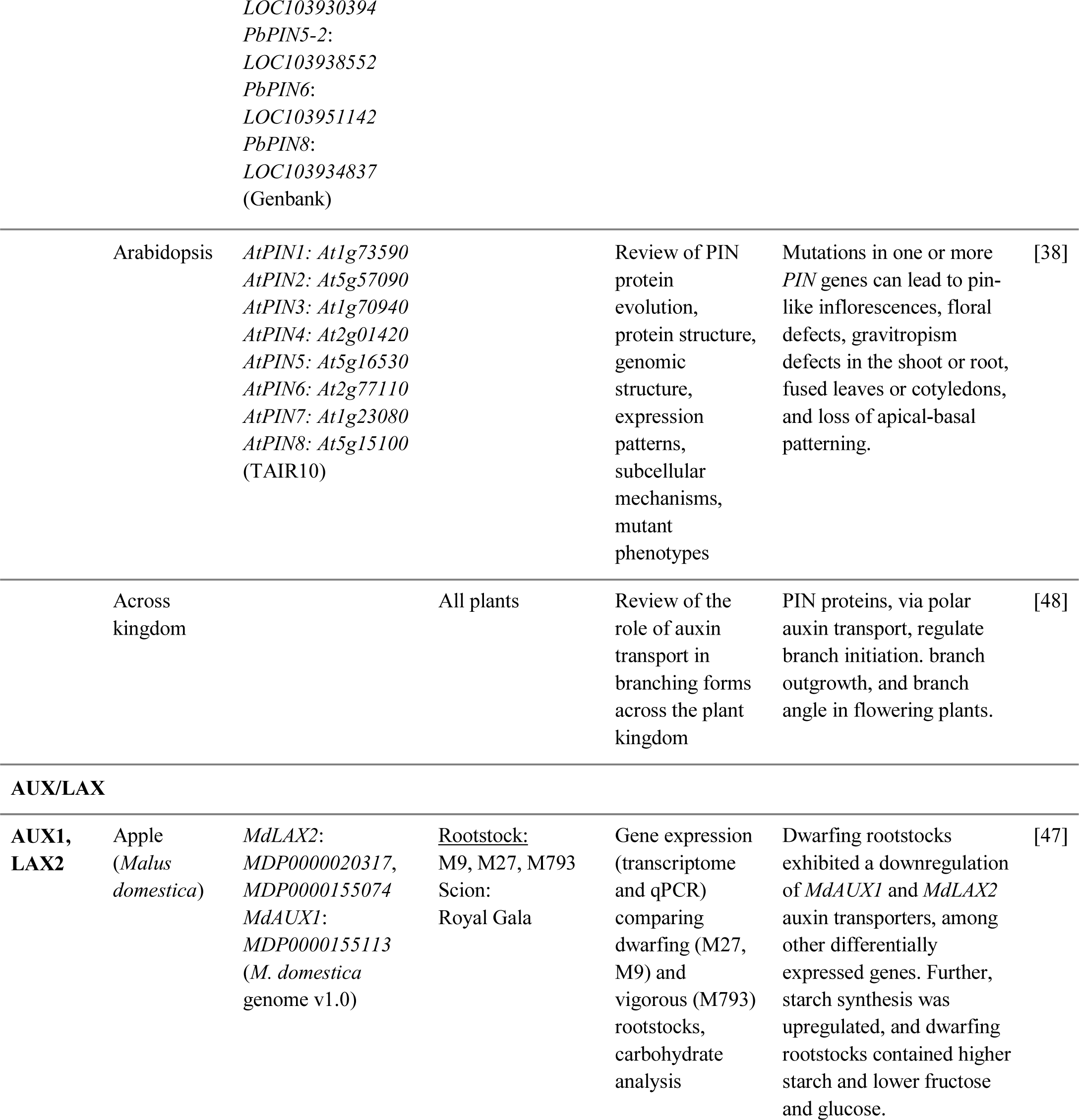
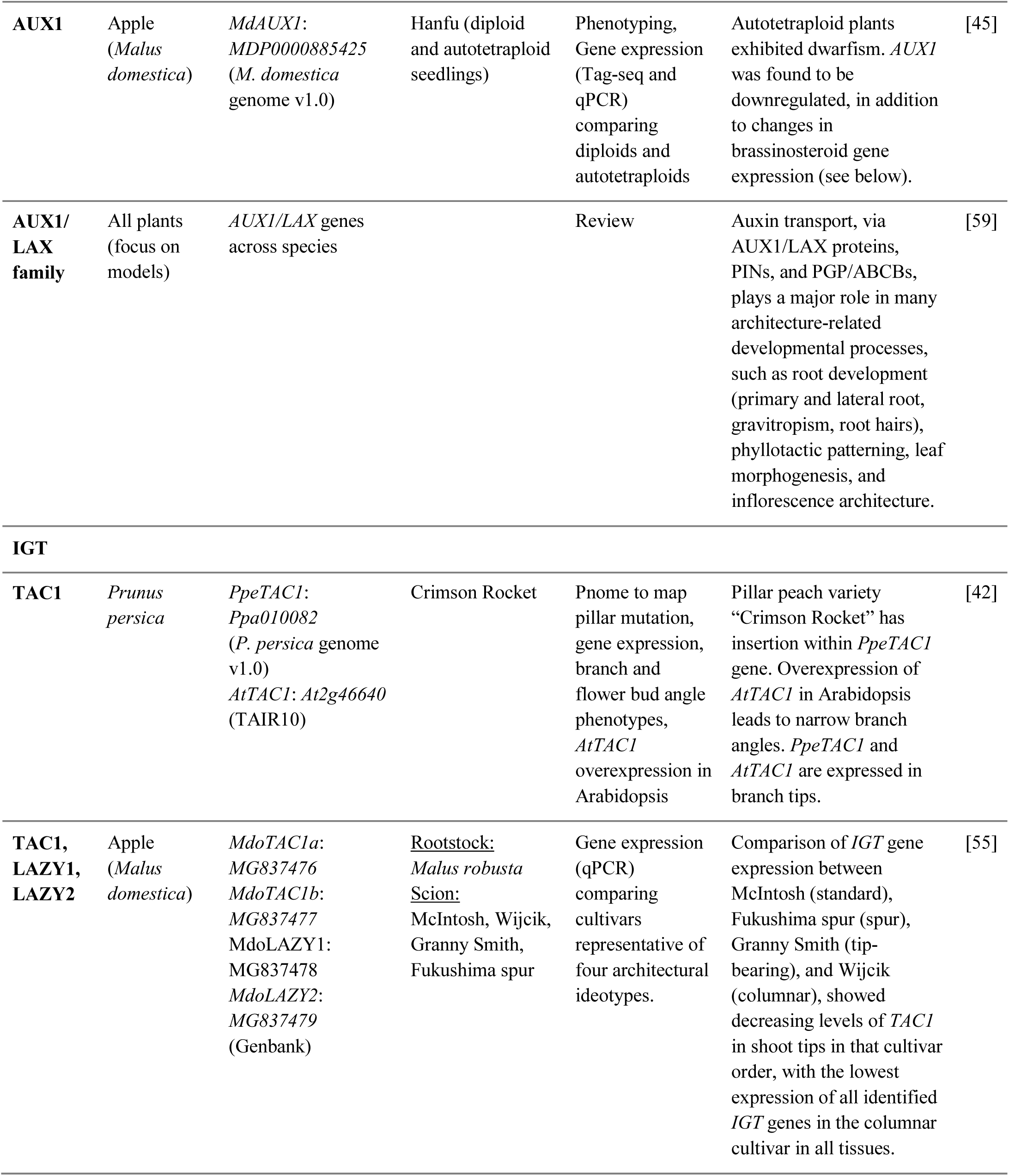
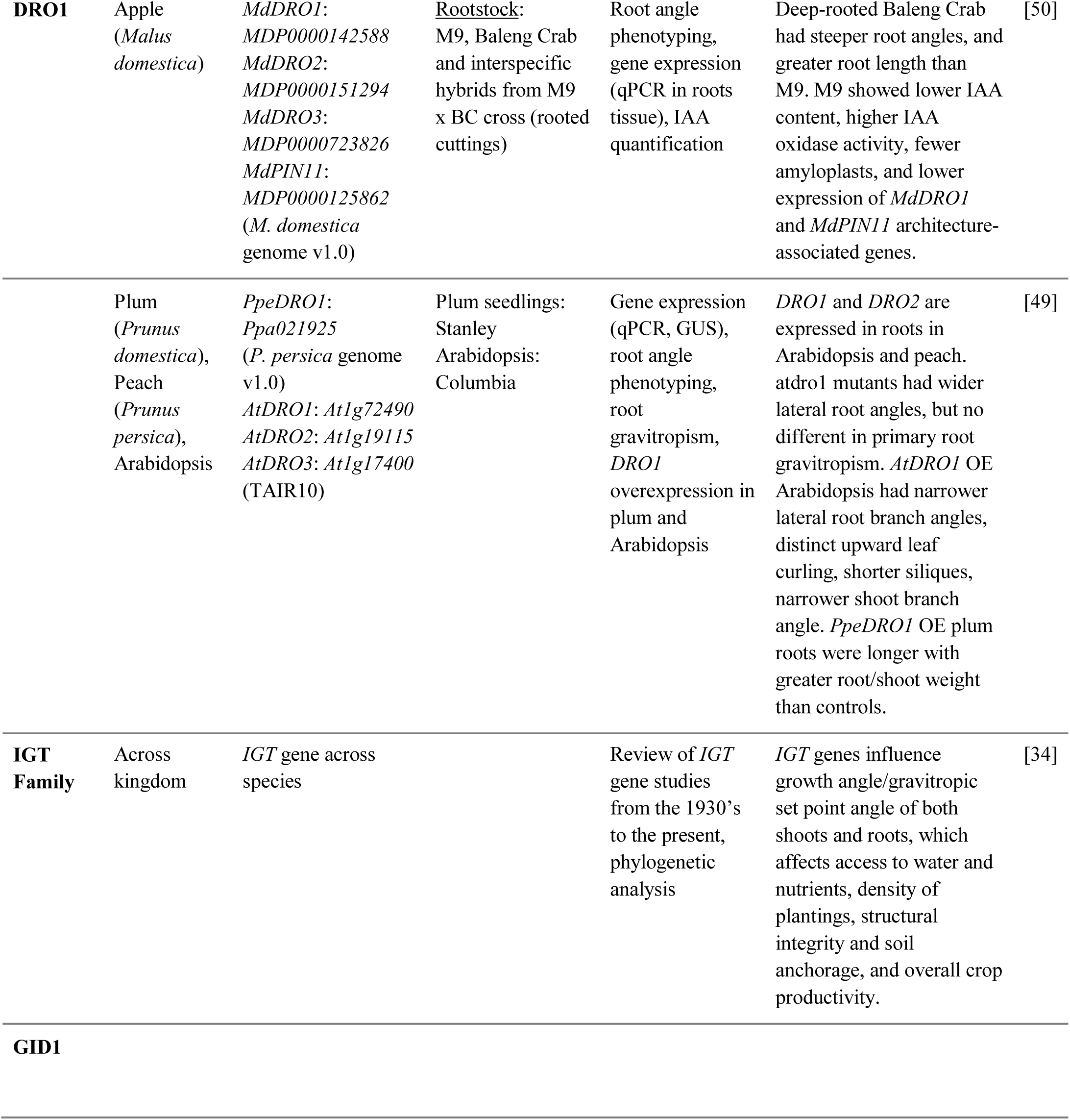
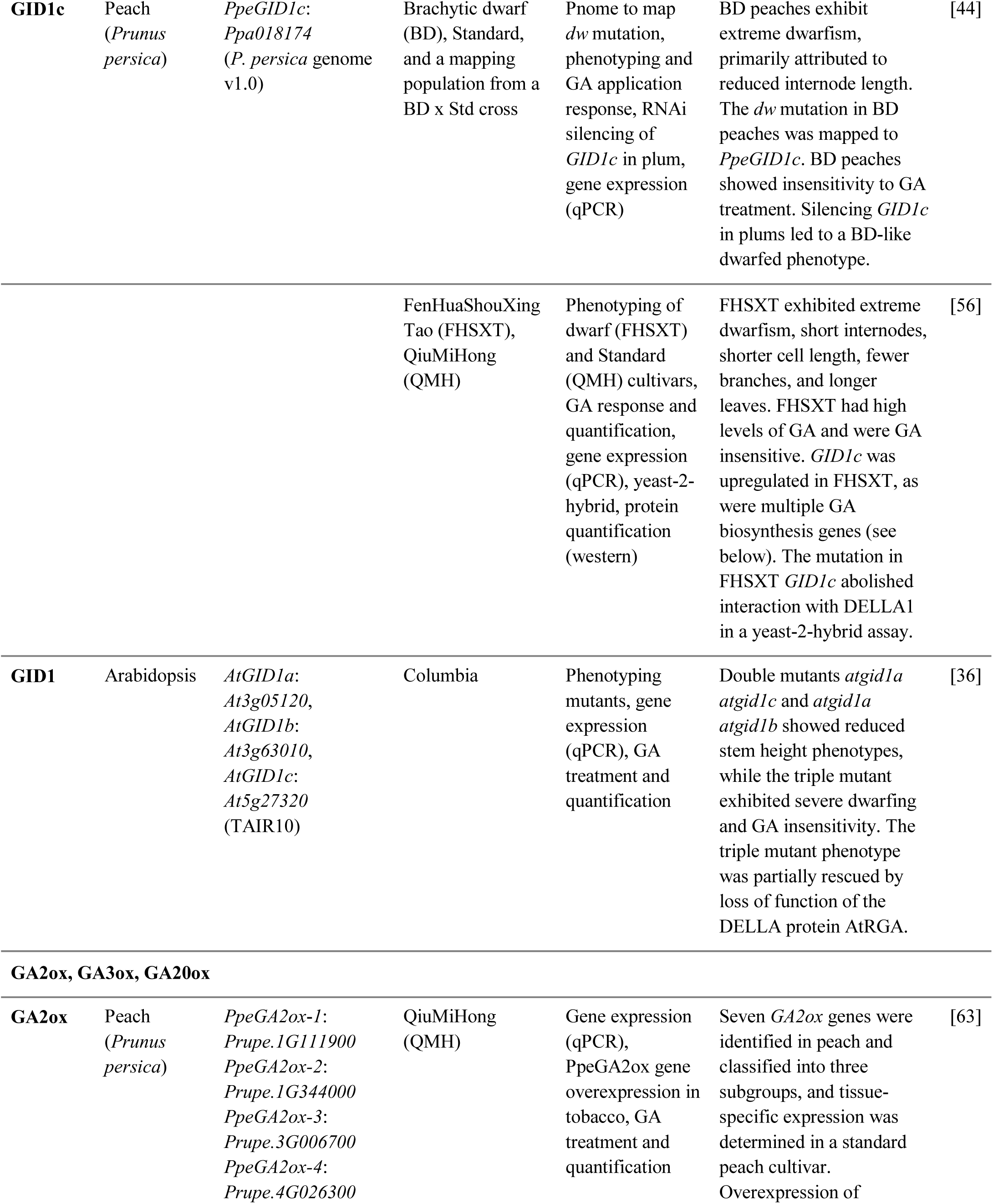
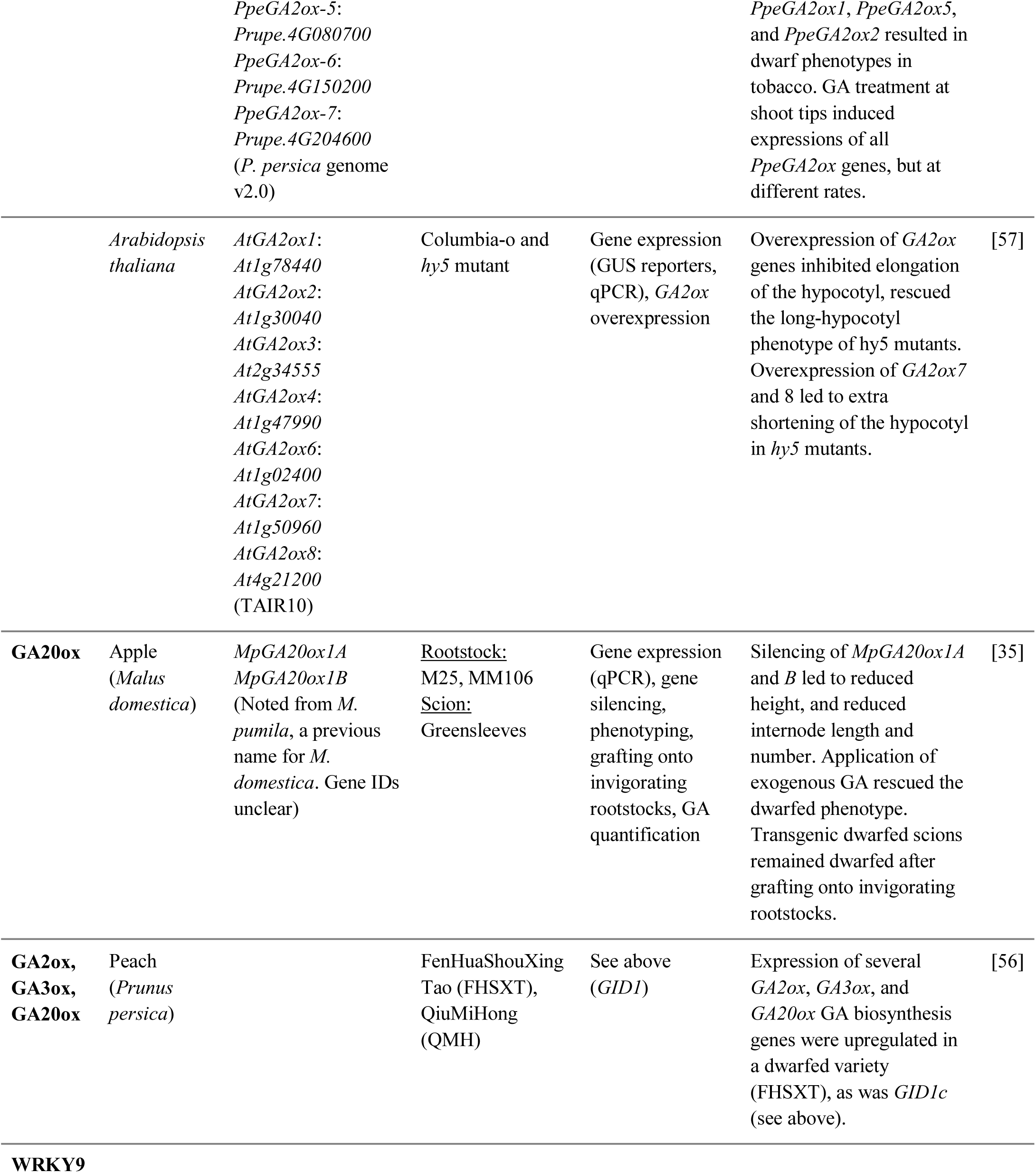
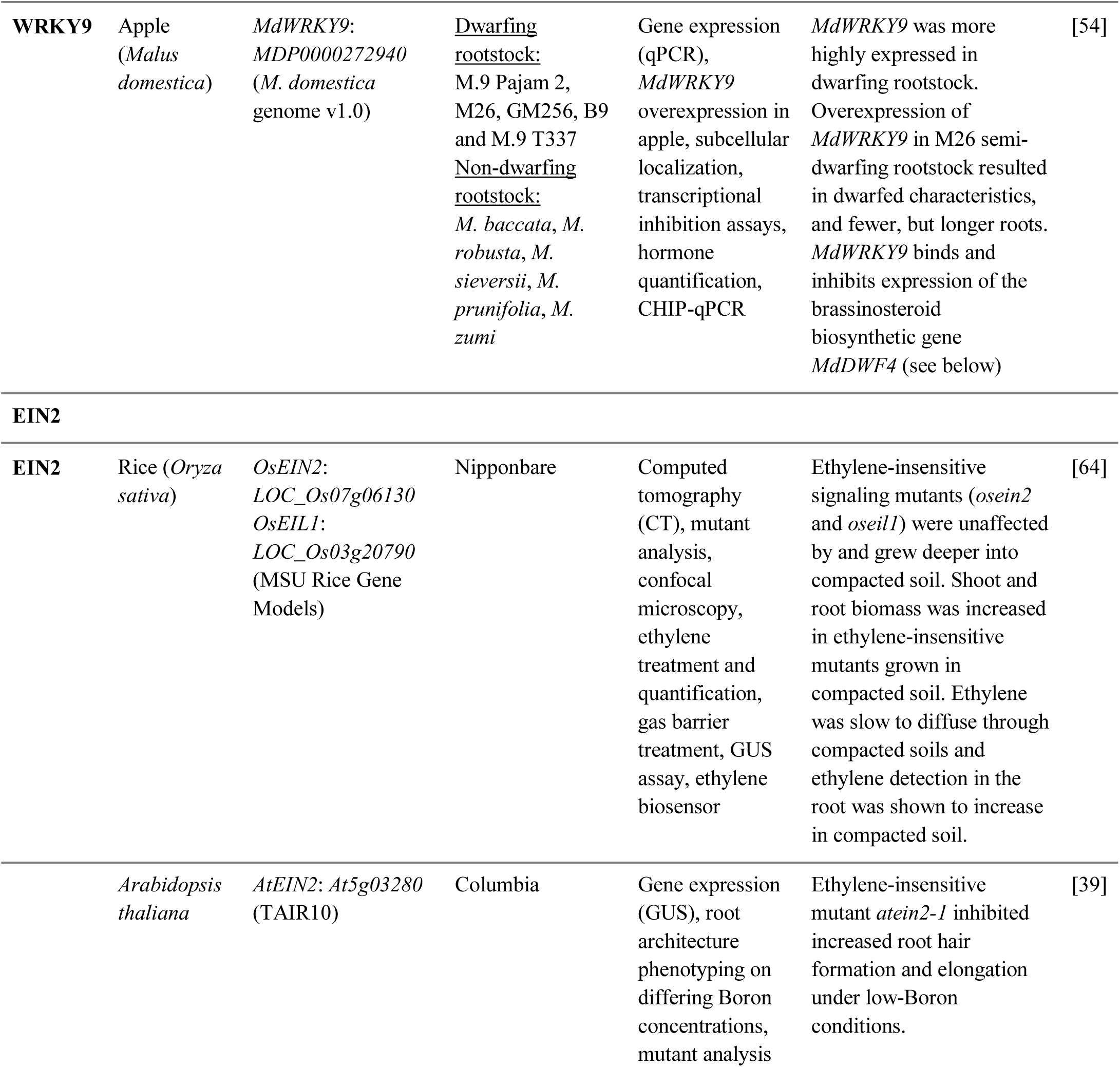
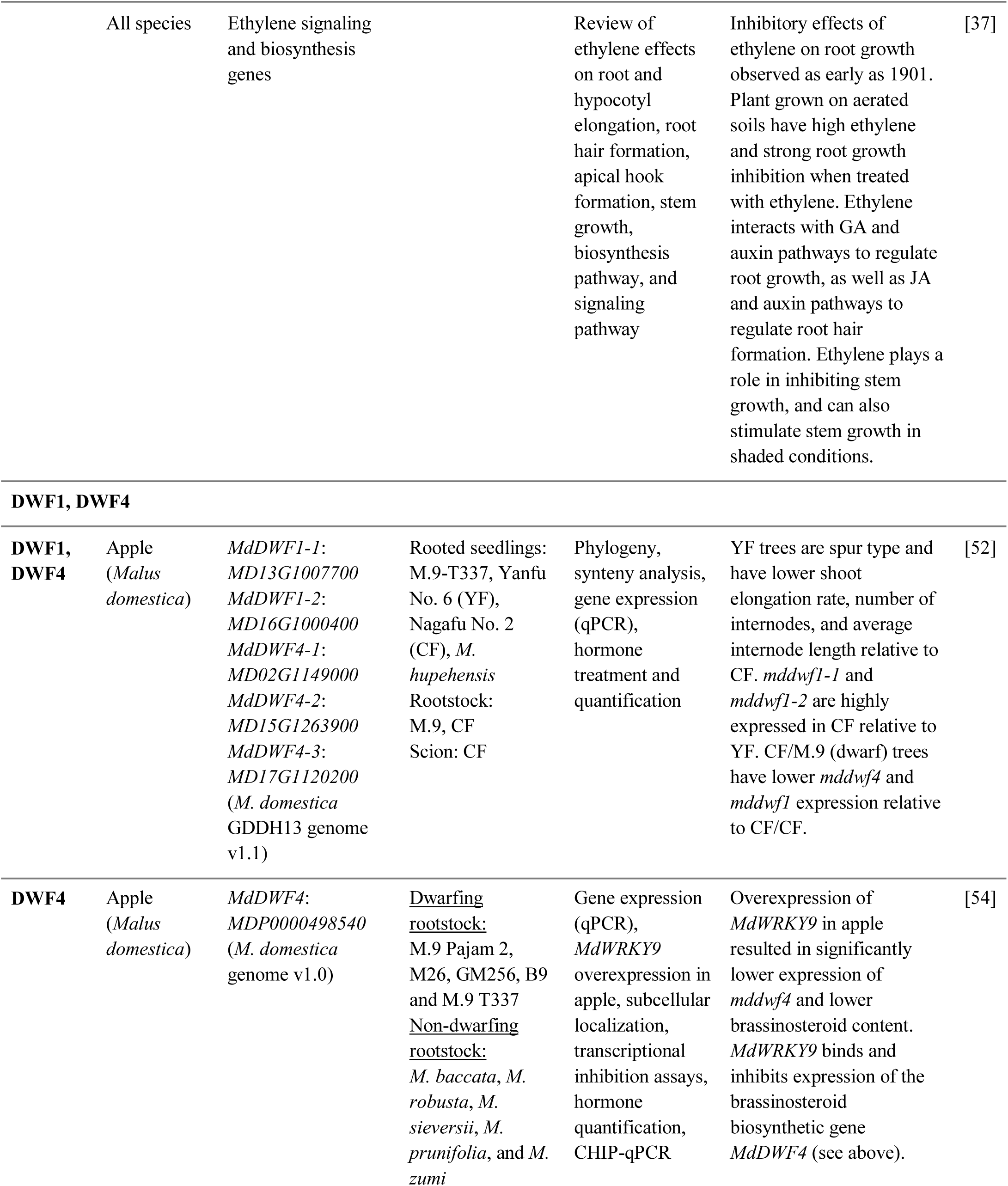
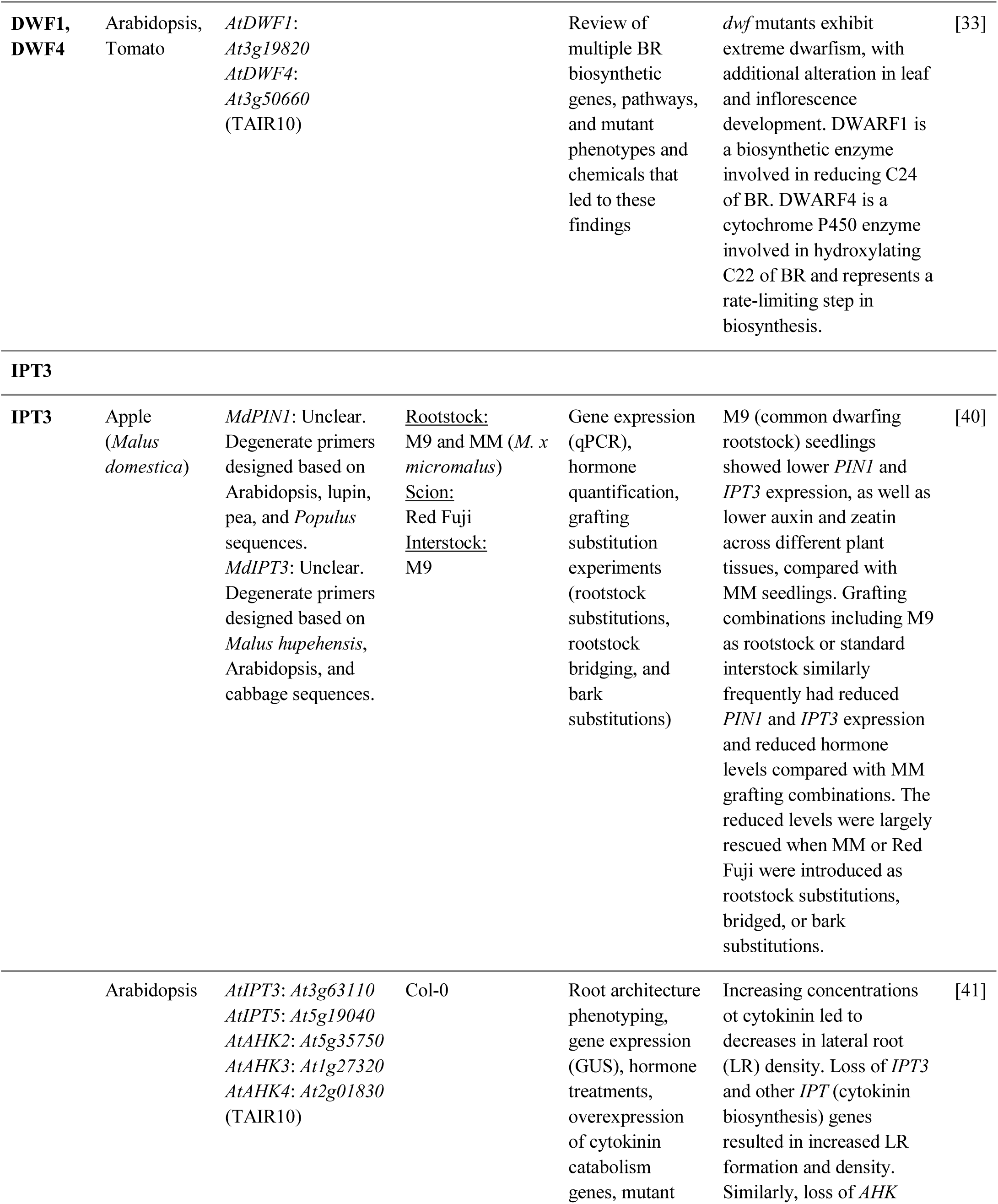
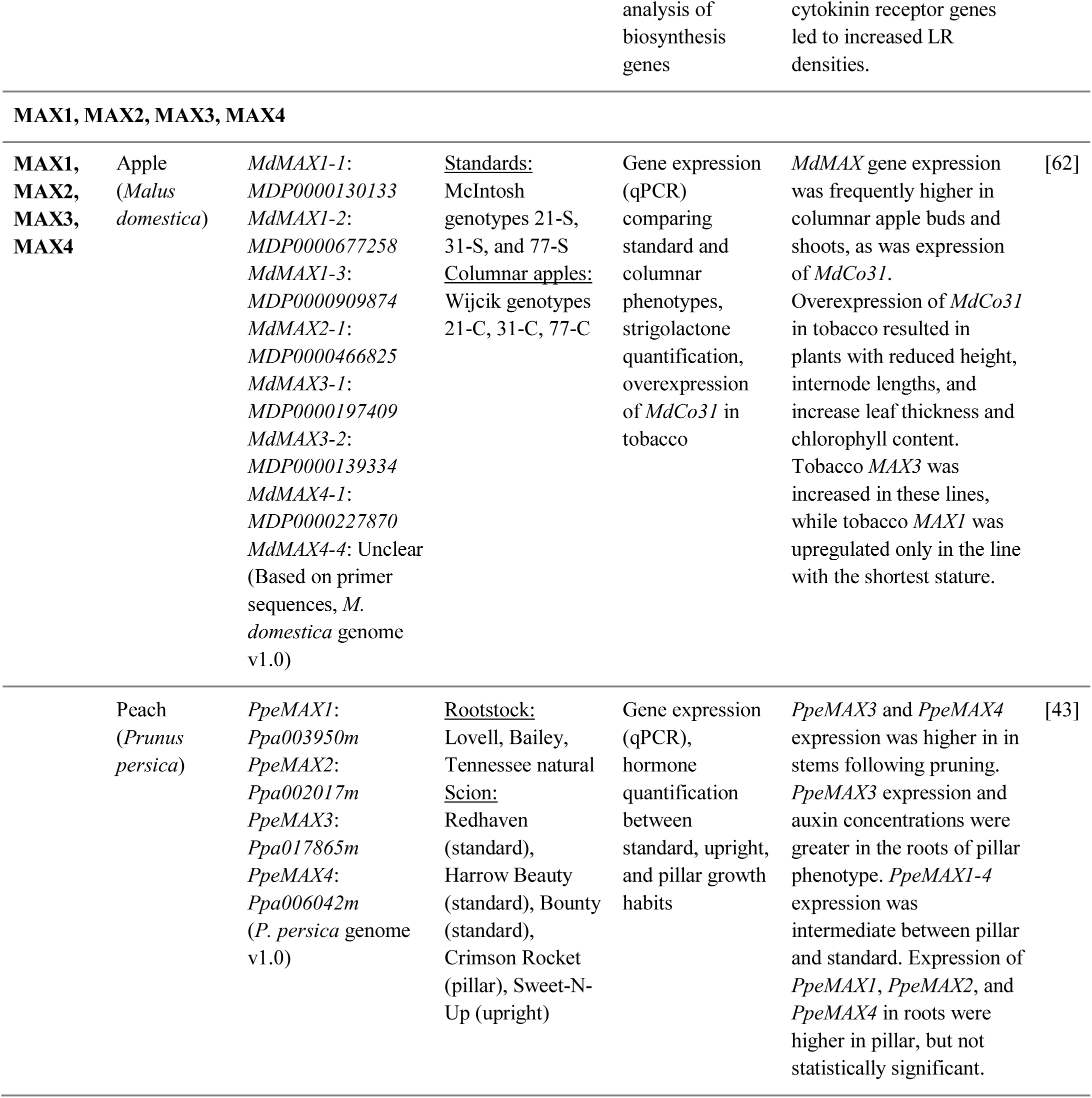
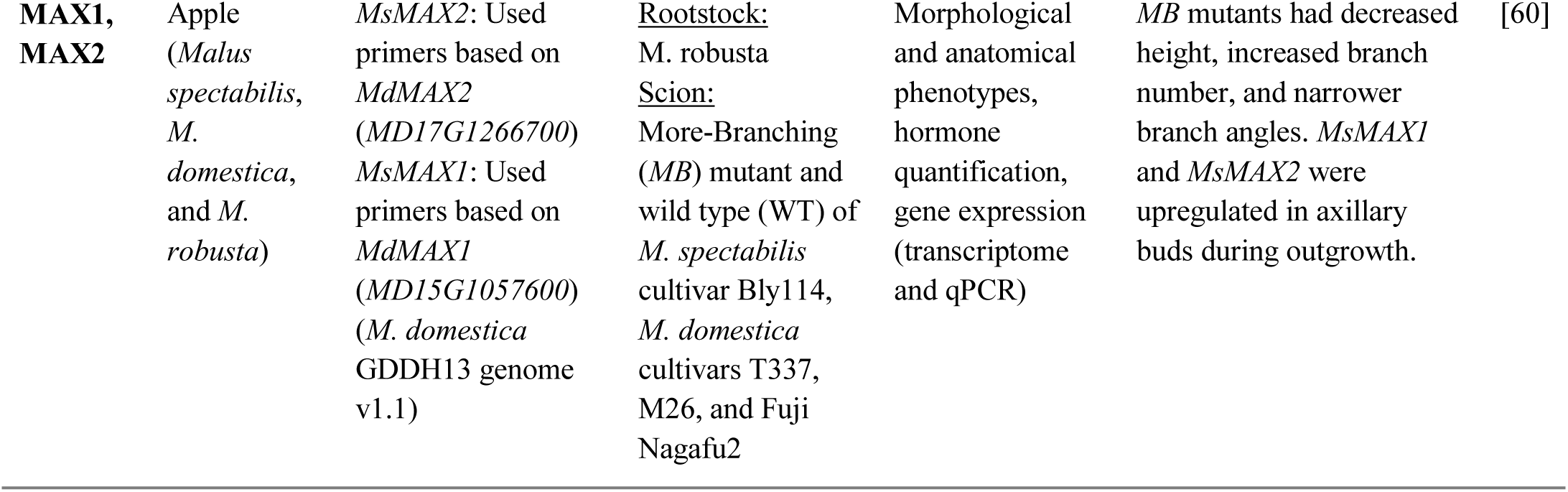
Architecture gene family table.

#### Overview of the gene identification workflow

Here, we developed a high throughput workflow (Fig. 2), leveraging a subset of the best Rosaceae plant genomes, and a phylogenomic perspective to efficiently and accurately generate lists of genes in gene families of interest and phylogenetic relationships of genes from different plant lineages. Our workflow, consisting of three main steps, implemented various functions from PlantTribes^65^ (https://github.com/dePamphilis/PlantTribes) and other software ^66, 67^ for targeted gene annotation.

**Fig 2.**
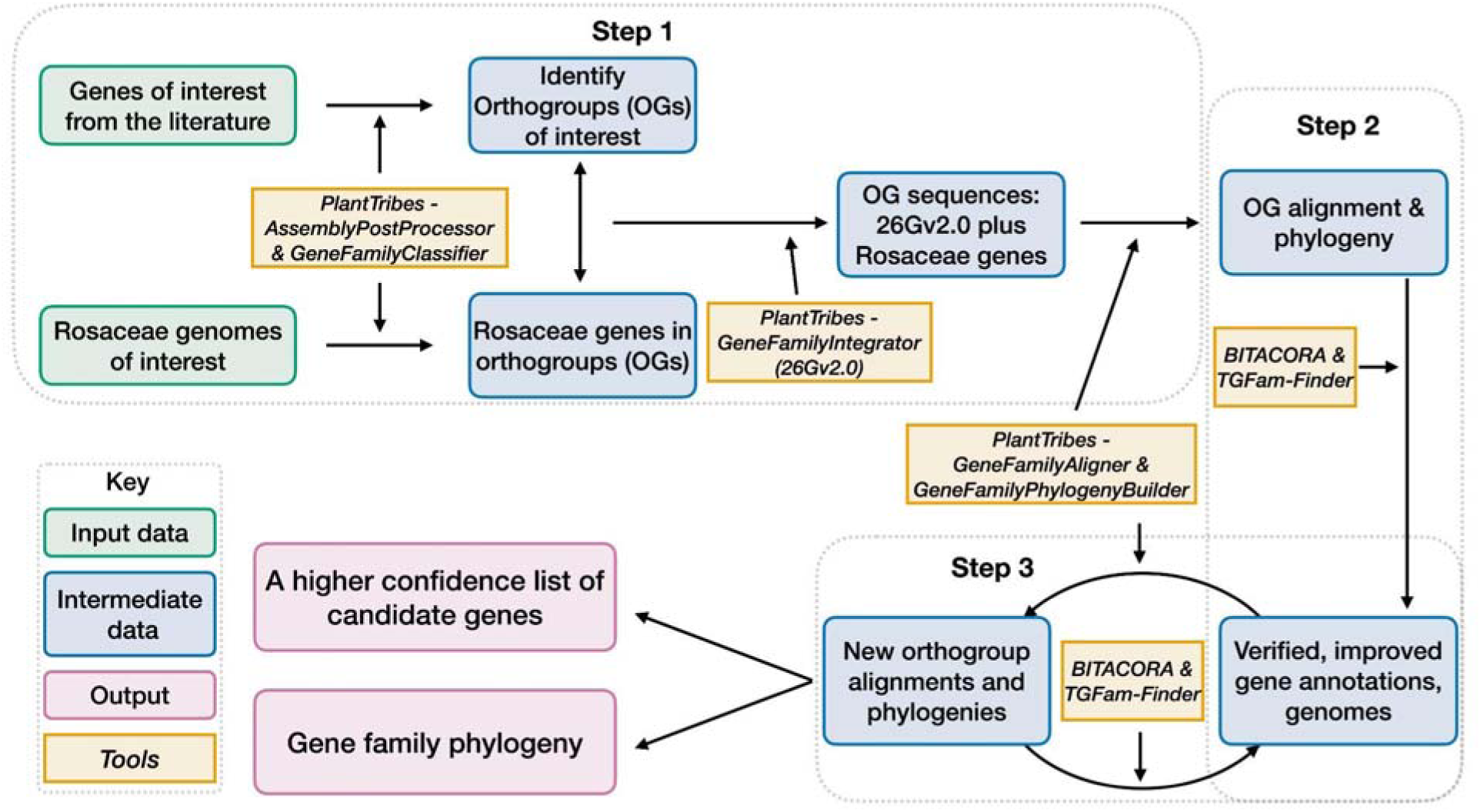
A workflow for candidate gene identification, curation, and gene family construction. Grey dotted boxes outlined the three steps of this workflow. Boxes with green outlines are input information. Boxes with blue outlines are intermediate outputs and boxes with purple outlines are final outputs. Contents in boxes with orange outlines are softwares used for generating the outputs.

#### Step 1 - An initial gene list and preliminary phylogenies

In Step 1, representative plant architecture genes obtained from the literature were assigned into orthogroups based on sequence similarity, giving us 22 orthogroups of interest (Supplementary Tables 4-5. Note that OG12636 is a monocot-specific orthogroup, thus not included in the downstream analysis of this section). In parallel, we classified all the genes annotated from 14 Rosaceae genomes (Supplementary Fig. 2) into the same database. Next, Rosaceae genes assigned into the 21 orthogroups were integrated with sequences from the 26 scaffolding species for multiple sequence alignments, which were used to infer phylogeny. At the end of this step, we obtained our initial list of genes in each orthogroup and the phylogenetic relationships of each gene family.

After examining the 21 orthogroups, we identified 64, 105, 94, and 53 genes from *Prunus persica*, Gala_v1, d’Anjou, and Bartlett.DH_v2, respectively. A whole genome duplication (WGD) event occurred in the common ancestor of *Malus* and *Pyrus*^14^, but was not shared with *Prunus.* Therefore, we expect to see an approximate 1:2 ratio in gene numbers in most cases, which explains fewer genes in *Prunus* compared to Gala_v1 and d’Anjou. However, the low gene count in Bartlett.DH_v2 was unexpected. For instance, we observed a clade within a PIN orthogroup (OG1145) comprised of short *PIN* genes^38^, which seemed to lack genes from the Bartlett.DH_v2 genome altogether (Fig. 3a). One gene copy is found in *Prunus* and Rosoideae species, and two copies are found in most of the Maleae species, but none were identified in Bartlett.DH_v2. In addition, in the four genomes mentioned above, we found a number of problematic genes (Supplementary Table 6), for example genes that appeared shorter than all other orthologs or contained unexpected indels likely due to assembly or annotation errors.

**Fig 3.**
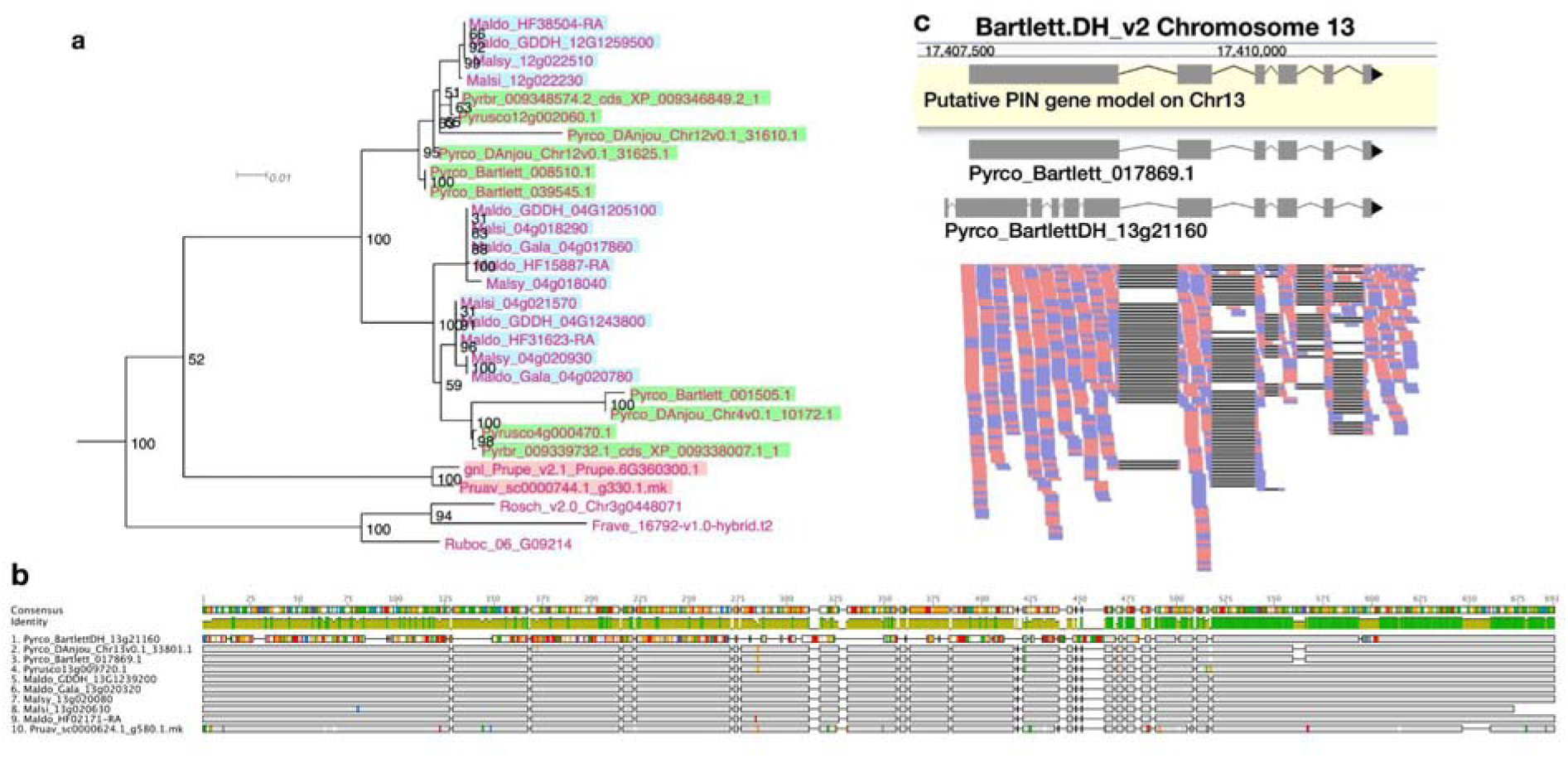
Phylogeny, amino acid sequence comparison, and RNAseq read mapping of *PIN* genes. a. One clade of short *PINs* from OG1145 phylogeny. *Malus* genes are indicated with a blue background, *Pyrus* with a green background, and *Prunus* with a pink background. **b.** Amino acid sequence alignment of orthologous genes from 10 Amygdaloideae species in the long *PIN* gene family (OG438). Sites identical to the consensus are shown in grey and sites different from the consensus are shown with a color following the Clustal color scheme in Geneious. Green color in the identity row indicates 100% identical across all sequences and greeny-brown color indicates identity from > 30% to < 100% identity. Gaps in the alignment are shown with a straight line. **c.** RNAseq reads (forward: red; reverse: blue) mapped to a fragment of chromosome 13 in the Bartlett.DH_v2 genome, where a long *PIN* gene, *Pyrco_BartlettDH_13g21160*, was annotated. Gene model in the yellow box is a putative gene model predicted with RNAseq reads (ref. 73) mapped to this region. The two gene models above the read mapping are retrieved from the original annotations of Bartlett_v1 (*Pyrco_Bartlett_017869.1*) and Bartlett.DH_v2 (*Pyrco_BartlettDH_13g21160*).

#### Step 2 and 3 - Iterative reannotation of problematic gene models

Inaccurate and missing gene models are common in any genome, especially in the early annotation versions^23, 24^. In model organisms, such as human, mouse (https://www.gencodegenes.org/), and *Arabidopsis* (https://www.arabidopsis.org/), gene annotations are continuously being improved using experimental evidence, improved data types (*e.g.* full-length RNA molecule sequencing), and both manual and computational curation. Building a better genome assembly is another way to detect additional genes. For instance, the BUSCO completeness score increased from 86.7% in the initial ‘Golden Delicious’ apple genome^16^ to 94.9% in the higher-quality GDDH13 genome^15^, indicating that the latter genome captured approximately 120 more conserved single-copy genes. Hence, we hypothesized that the potentially missing and problematic gene models we observed in the two European pears could be improved by: 1) using additional gene annotation approaches; and 2) searching against improved genome assemblies.

To test whether further gene annotation would improve problematic gene models, we moved forward to Step 2 of our workflow, using results from Step 1 as inputs. For each orthogroup containing problematic European pear genes (Supplementary Table 6), we used a subset of high-quality gene models from Rosids identified in Step 1 as inputs and re-annotated these gene families in the two pear genomes. After using a combination of annotation softwares and manual curation, we found a total of 98 genes from the d’Anjou genome, and reduced the number of problematic or incomplete genes from 34 to 3. In Bartlett.DH_v2, we identified 20 complete genes that were not annotated in the original genome and improved the sequences of 7 previously problematic genes, however, the total number of the selected architecture genes (73 genes among which 15 were problematic or incomplete) was still notably lower than that of d’Anjou (98 with 3 incomplete genes) or Gala (105 with 15 being incomplete, see Supplementary Table 6). In Step 3, which involves iterative steps of phylogenetic analysis and targeted gene re-annotation, we added additional information such as the improved d’Anjou genes and RNA-seq datasets as new resources to annotate Bartlett.DH_v2 genes, but found no improvements in identifying unannotated genes or improving problematic models.

Results gathered after the first iteration of Step 3 supported our hypothesis that extra annotation steps could help improve imperfect gene models and identify missing genes in the two targeted European pear genomes. However, there were still about 30 genes potentially missing in Bartlett.DH_v2, which led us to test whether polishing the genome assembly would further improve problematic or missing gene models.

#### Step 3 - Adding Bartlett.DH_v2 genome polishing

The quality of genome assembly is affected by many factors, including sequencing depth, contig contiguity, and post-assembly polishing. Attempts to improve a presumably high-quality genome are time consuming, and may prove useless if the genome is already in good condition. To initially determine whether polishing the genome assembly would be useful, we first investigated the orthogroups with problematic Bartlett.DH_v2 genes to seek for evidence of assembly derived annotation issues. Indeed, in most cases where we failed to annotate a gene from presumably the correct genomic region, we observed unexpected indels while comparing the Bartlett.DH_v2 genome assembly to other pear genomes (Supplementary Fig. 3 and Supplementary Table 7).

Unexpected indels in the Bartlett.DH_v2 genome were associated with incorrect gene models as well. For example, Fig. 3b shows a subset of amino acid sequence alignments for a specific member (*Pyrco_BartlettDH_13g21160*) of a PIN orthogroup (OG438) comprised of the long *PIN* genes^38^, in which the Bartlett.DH_v2 gene model shared low sequence identity with orthologs from other Maleae species and *Prunus*. To validate the identity of the problematic gene models, we leveraged RNAseq data from various resources^68–73^ and mapped them to the Bartlett.DH_v2 gene models. In most cases where a conflict was present between the pear consensus, for a given gene of interest, and the Bartlett.DH_v2 gene model, the reads supported the consensus (Fig. 3c). The frequent occurrence of truncated and missing genes in the Bartlett.DH_v2 genome may be caused by assembly errors (*e.g.,* base call errors, adapter contamination) that create erroneous open reading frames. This observation provided us with the first piece of evidence that the differences in gene family content observed in the Bartlett.DH_v2 genome may not only be caused by misannotations, but also assembly issues.

To further test whether improvement to the genome assembly would allow us to capture the problematic and missing genes, we polished the Bartlett.DH_v2 genome with Illumina reads from the original publication ^17^. We identified 98.40% complete BUSCOs in the polished genome assembly, a 1.90% increase compared to the original assembly (Supplementary Table 2).

Using the polished genome, we reiterated Step 3 of our workflow and annotated a total of 103 genes in our gene families of interest, with only two gene models being incomplete (Supplementary Table 6). This new result doubled the number of genes we identified from the original genome annotation and brought the expected gene number into parity with other pome fruit genomes. This supports our hypothesis that genes were missing due to methodological reasons, and in this case, due to assembly errors.

### Curation of a challenging gene family: the IGT family

Some gene families are more complex than others. For example, it is more difficult to study the evolution of resistance (R) genes than most BUSCO genes because the former is comprised of fast-evolving multigene families while the latter are universally conserved single-copy gene families. Within the architecture gene families we studied, the IGT family is more challenging than many others because members of this family have relatively low levels of sequence conservation outside of a few conserved domains^74^. Previous reports identified four major clades (LAZY1-like, DRO1-like, TAC1-like, and LAZY5-like) in this gene family^34^. Study of LAZY1 in model species identified 5 conserved regions^74^ (Fig 4c). The same domains are also present in other LAZY1-like and DRO1-like proteins and the first 4 domains are found in TAC1-like proteins across land plants^75^. LAZY5-like, the function of which is largely unknown, has only domains I and V. Early research of the *TAC1-like* and *LAZY1-like IGT* genes identified these genes as grass-specific^76, 77^, as BLAST searches failed to find homologs in other plant lineages.

**Fig 4.**
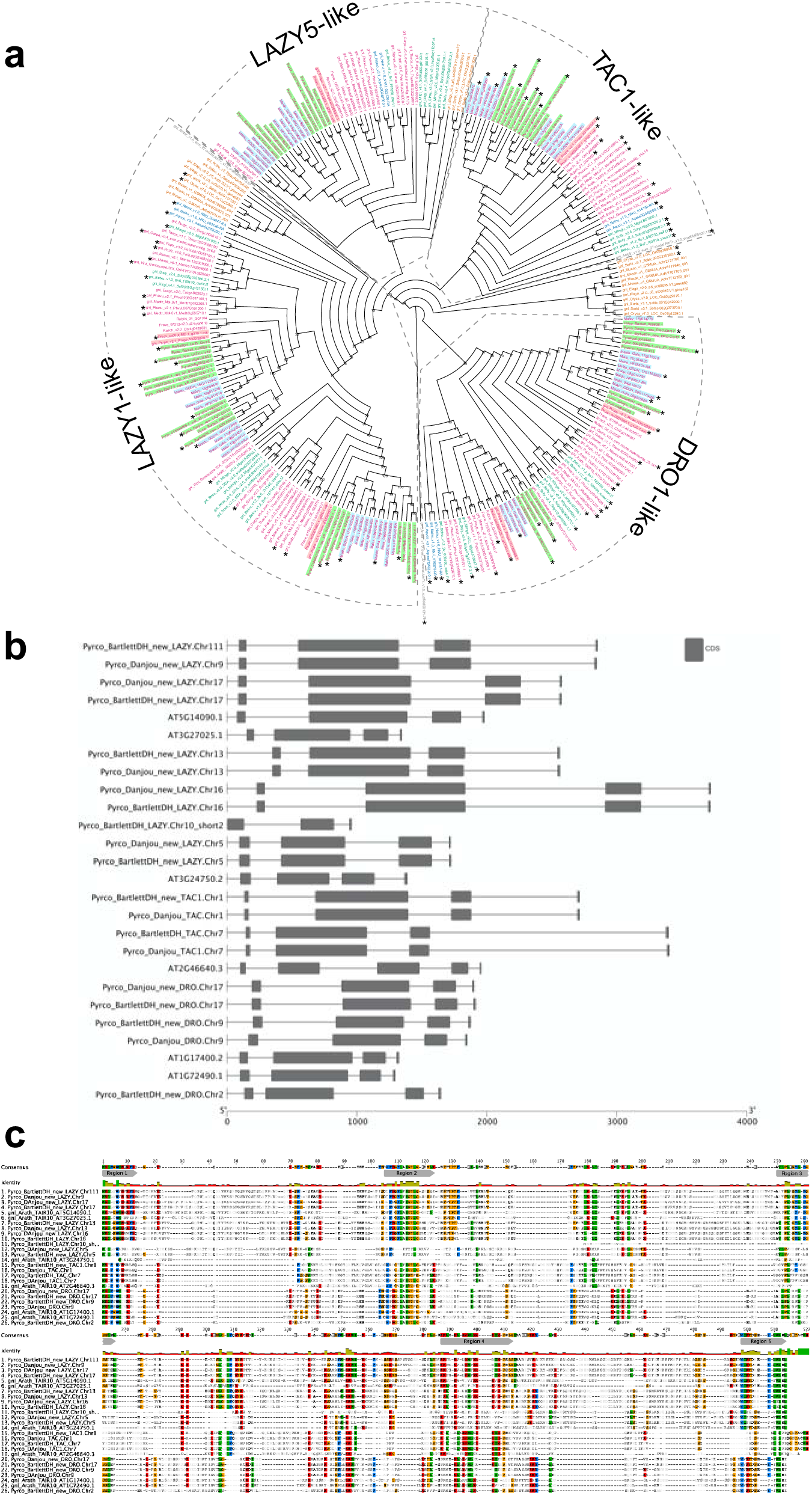
Phylogeny, intron-exon structure, and amino acid comparison of IGT genes. a. Cladogram of the IGT gene family (including LAZY1-like, LAZY5-like, TAC1-like, and DRO-like). Genes are colored as shown in Supplementary Fig. 2. 1000 bootstrap replicates were conducted to estimate reliability and the numbers on the node indicate bootstrap support. **b.** Cartoon illustrating intron-exon structures of IGT genes from *Arabidopsis thaliana* (Araport11), Bartlett.DH_v2, and d’Anjou. **c.** Amino acid alignment of IGT genes from *Arabidopsis thaliana*, Bartlett.DH_v2, and d’Anjou. Sites consisting of a similar amino acid type as the consensus were highlighted with a background color following the Clustal color scheme in Geneious. Red color in the identity row indicates identity < 30%. Five conserved regions were highlighted with a grey symbol below the consensus sequence.

Using Arabidopsis and rice *IGT* genes as queries, our workflow identified five orthogroups (Supplementary Table 4), containing all the pre-characterized *IGT* genes in angiosperms. The phylogeny constructed with these five orthogroups largely supported previous classification of the four clades^34^, and provided more information regarding the evolutionary history of this gene family (Fig. 4a and Supplementary Fig. 4). The TAC1-like clade, which is sister to the others, is divided into two monophyletic groups; one contains only monocots while the other has representatives from all the other angiosperm lineages. The LAZY1-like and LAZY5-like clades form one large monophyletic group, which is sister to the DRO1-like clade. Within Rosaceae, a near 1:2 ratio was expected between peach and pear due to the WGD in the common ancestor of the Maleaes. Compared to the six known peach *IGT* genes^34^, we found 11 orthologs in Bartlett.DH_v2 (including 1 short gene, *Pycro_BartlettDH_LAZY.Chr10*, caused by an unexpected premature stop codon) and 9 in d’Anjou (*Pycro_Danjou_DRO.Chr2* and *Pycro_Danjou_LAZY.Chr10* failed to be annotated due to missing information in the genome). The resulting phylogeny (Fig. 4a) shows that we have now identified most of the expected *IGT* genes in European pears.

Besides low sequence similarity, *IGT* genes also have unique intron-exon arrangements, which are conserved across Arabidopsis and a few other plant species^34, 74, 78^. These genes all contain 5 exons, but unlike most genes, the first exon only comprises six nucleotides and the last exon contains ∼20 nucleotides. Annotation of short exons, especially when transcriptome evidence is limited, can be very challenging and skipping such exons could cause problems in gene discovery^79–81^. For instance, the annotation of *AtAPC11* (*At3g05870*) was inaccurate until Guo and Liu identified a single-nucleotide exon in this gene^80^.

To determine whether we captured the correct *IGT* gene models in the targeted genomes, we investigated the protein sequence alignments and gene features. In the original annotation, only three gene models (*Pyrco_BartlettDH_16g10510*, *Pyrco_BartlettDH_07g15250*, *Pyrco_DAnjou_Chr7v0.1_17442.1*) have the correct intron-exon combination and the expected domains. In the iterative re-annotation steps of our workflow, we identified 6 additional accurate gene models leveraging sequence orthology and transcriptome evidence^68–73^. We further investigated all the sequences we identified as *IGT* genes, seeking the presence or absence of the expected domain features. However, even among gene models from the best annotated genomes used to construct the 26Gv2.0 database, only 45.16% (56/124) have the expected domain features (Indicated with an * next to gene names in Fig. 4a) (LAZY5-like was not taken into consideration due to its unique structure). In most cases, although the signature IGT domain (II) is correctly identified in the genes, domains I and V are usually missing or incorrect, likely due to mis-annotation of the first and last short exons. In Rosaceae, besides Bartlett.DH_v2 and d’Anjou, 34.38% (33/96) had the expected domains (Fig. 4a). This finding motivated us to manually investigate the targeted genomes to annotate the IGT genes. Using the correct gene models as reference, plus a careful manual curation, we were able to annotate 19 complete gene models of 20 expected IGT genes from the two targeted pear genomes (Figs. 4b and 4c).

## Discussion

A second European pear cultivar genome from ‘d’Anjou’ provided additional insights into gene families across Rosaceae. By leveraging perspectives from comparative genomics and phylogenomics, we developed a high-throughput workflow using a collection of bioinformatic tools that takes a list of genes of interest from the literature and genomes of interest as input, and produces a curated list of the targeted genes in the query genomes.

In the case study presented here, candidate genes from 16 plant architecture-related gene families were identified from 14 Rosaceae genomes. The study of gene families consists primarily of two initial parts: first, identification of all the members in these families, and second, investigation of their phylogenetic relationships. Many attempts^82–84^ to identify genes of interest from a genome have relied solely on a BLAST search querying a homolog from a model organism, which may be distantly related. However, such a method is insufficient in identifying all members of a large complex gene family or a fast-evolving and highly-divergent family, such as the *IGT* genes.

They may also incorrectly include genes in a gene family based only on one or a few highly conserved regions that are insufficient for gene family membership. Compared to a BLAST-only approach, the gene classification process in our workflow used a combination of BLAST and HMMER search of an objectively pre-classified gene family scaffold, which provides a better result by taking into consideration both sensitivity and specificity^65^. This allowed us to efficiently identify even very challenging genes. Moreover, phylogenetic relationships revealed by a small number of taxa, for instance using only one species of interest and one model organism, can be inaccurate. For example, in our phylogenetic analysis with rich taxon sampling, *PIN5-1* and *PIN5-2* from *Pyrus bretschneideri* are sisters to all other *PINs* (Supplementary Fig. 5), challenging the phylogenetic relationship inferred with *PINs* only from *P. bretschneideri* and *Arabidopsis thaliana*^61^.

The iterative quality control steps in the workflow helped identify problems that existed in certain gene models and provided hints about where to make targeted improvements to important *Pyrus* genomic resources. The highly contiguous assembly of Bartlett.DH_v2 provided a valuable reference to anchor the shorter scaffolds from d’Anjou, which is essential for a good annotation. On the other hand, the perspective afforded by the d’Anjou genome led us to examine the Bartlett.DH_v2 genome assembly further. We developed and tested hypotheses regarding unexpected gene annotation patterns in the two targeted European pear genomes among various Maleae species and cultivars. This led to a polished assembly and improved annotations that allowed us to curate a high confidence list of candidate genes and gene models for downstream analyses. By adding targeted iterations of genome assembly and annotation, we now have a better starting point for reverse genetic analyses and understanding functionality of architecture-related genes in pears.

The challenges we encountered as we laid the groundwork for reverse genetics studies to understand pear architecture genes, and the approaches we took to evaluate and tackle these challenges, reinforce the idea that genome assembly and annotation are iterative processes. We found that relating gene accession IDs and inconsistent gene names back to gene sequences in various databases was often difficult and time consuming. Objective, global-scale gene classification, as we used here via PlantTribes^65, 85^, can help researchers work across genomes and among various genome resources. Further, guidance from consortia such as AgBioData^86^ is helping facilitate work such as we have described here that includes the acquisition and analysis of genome-scale data. Our starting point for understanding putative architecture genes in pear was with genes of interest from several plant species - an approach that many researchers will find familiar. With genes of interest in hand, our workflow provides a comparative genome approach to efficiently identify, investigate, and then improve and/or validate genes of interest across genomes and genome resources.

## Materials and Methods

### Plant materials and sequencing

The ‘d’Anjou’ plants were purchased from Van Well’s nursery in East Wenatchee, WA, USA and grown in the USDA ARS greenhouse #6 at Wenatchee, WA, USA. Fresh leaves (∼1.5g) from one ‘d’Anjou’ plant were flash frozen and used for DNA extraction. A CTAB isolation protocol^87^ was used to generate high-molecular-weight genomic DNA with the following modifications: the extraction was performed large scale in 100 ml of extraction buffer in a 250 ml Nalgene centrifuge bottle; the isopropanol precipitation was performed at room temperature (∼ 5 minutes) followed immediately by centrifugation; after a 15-minute incubation in the first pellet wash solution, the pellet was transferred to a 50 ml centrifugation tube via sterile glass hook before performing the second pellet wash; following the second pellet wash, centrifugation and air drying, the pellet was resuspended in 2 ml TE buffer (10 mM Tris, 1 mM EDTA, pH 8.0) and allowed to resuspend at 4 °C overnight. The concentration of the DNA was measured by a Qubit 2.0 fluorometer (Invitrogen) and 50 ug DNA was digested with RNase A (Qiagen, final concentration 10 ug/ml, 37 °C for 30 minutes) and then further cleaned up using the PacBio recommended, user-shared gDNA clean-up protocol (https://www.pacb.com/search/?q=user+shared+protocols) performed at large-scale with the DNA sample brought up to 2 ml with TE and all other volumes scaled up accordingly. The final pellet was resuspended in 100 ul TE. The final DNA concentration was measured by Qubit fluorometer, and 500 ng was loaded onto a PFG (Bio-Rad CHEF) to check the size range. The DNA ranged in size from 15 Kb to 100 Kb with a mean fragment size around 50 Kb. The purity of the DNA as measured by the NanoDrop spectrophotometer (ThermoFisher) was 260/280 nm: 1.91; 260/230 nm: 2.51. Cleaned-up gDNA was sent to the Penn State Genomics Core facility (University Park, PA, USA) for Pacbio and Illumina library construction and sequencing. A total of 10 ug gDNA was used to construct PacBio SMRTbell libraries and sequenced on a PacBio Sequel system. A small subset of the same gDNA was used to make Illumina TruSeq library and was sequenced on an Illumina HiSeq 2500 platform. In addition, 4 ug of the same gDNA was sent to the DNA technologies and Expression Analysis Core Laboratory at UC Davis (Davis, CA, USA) to construct an Illumina 10X Chromium library, which was sequenced on an Illumina NovaSeq 6000 sequencer.

### Genome assembly and post-assembly processing

To create the initial backbone assembly of d’Anjou, Canu assembler v2.1.1^88^ was used to correct and trim PacBio continuous long reads (CLR) followed by a hybrid assembly of Illumina short reads and PacBio CLR with MaSuRCA assembler v3.3.2^89^. Next, Supernova v2.1.1, the 10x Genomics *de novo* assembler^90^, was used to assemble linked-reads at five different raw read coverage depths of approximately 50x, 59x, 67x, 78x, and 83x based on the kmer estimated genome size, and the resulting phased assembly graph was translated to produce two parallel pseudo-haplotype sequence representations of the genome. The Supernova assembler can only handle raw data between 30- to 85-fold coverage of the estimated genome size. Therefore, the muti-coverage assemblies provide an opportunity to capture most of the genome represented in the ∼234-fold coverage sequenced 10x Chromium read data. One of the pseudo-haplotypes at each of the five coverages was utilized for subsequent meta-assembly construction to improve the backbone assembly using a combination of assembly metrics, including 1) contig and scaffold contiguity (L50), 2) completeness of annotated conserved land plants (embryophyta) single-copy BUSCO genes^30^, and 3) an assembly size closer to the expected d’Anjou haploid genome size. The backbone assembly was incrementally improved by bridging gaps and joining contigs with the Quickmerge program^91^ using contigs from the five primary Supernova assemblies in decreasing order of assembly quality. The resulting meta-assembly at each merging step was only retained if improvement in contiguity, completeness, and assembly size was observed.

Next, the long-distance information of DNA molecules provided in linked-reads was used to correct assembly errors introduced in the meta-assembly during both the *de novo* and merging steps of the assembly process with Tigmint^92^ and ARCS^93^ algorithms. Tigmint aligns linked reads to an assembly to identify potential errors and breaks assembled sequences at the boundaries of these errors. The assembly is then re-scaffolded into highly contiguous sequences with ARCS utilizing the long-distance information contained in the linked reads. To further improve the d’Anjou meta-assembly, trimmed paired-reads from both the short insert Illumina and 10x Chromium libraries were utilized to iteratively fill gaps between contigs using GapFiller v1.10^94^, and correct base errors and local misassemblies with Pilon v1.23^95^. The genome assembly process is illustrated in Supplementary Fig. 6.

### Pseudomolecule construction

Before constructing the d’Anjou nuclear chromosomal-scale pseudomolecules, extraneous DNA sequences present in meta-assembly were identified and excluded (Supplementary Fig. 6).

Megablast searches with e-value < 1e-10 was performed against the NCBI nucleotide collection database (nt), and then the best matching Megablast hits (max_target_seqs = 100) against the NCBI taxonomy database were queried to determine their taxonomic attributions. Assembly sequences with all their best-matching sequences not classified as embryophytes (land plants) were considered contaminants and discarded. A second iteration of Megablast searches of all the remaining sequences (embryophytes) was performed against the NCBI RefSeq plant organelles database to identify chloroplast and mitochondrion sequences and assembly sequences with high similarity (> 80% identity; > 50% coverage) to plant organelle sequences were discarded ^27, 96^. Finally, the remaining meta-assembly nuclear contigs and scaffolds were ordered and oriented into chromosomal-scale pseudomolecules with RaGOO^97^ using the European pear, *Pyrus communis* Bartlett.DH_v2 genome^17^ reference chromosomes (Supplementary Fig. 6).

### Assembly validation

Both the contig and scaffold assembly metrics were evaluated in addition to the completeness of universally conserved single-copy genes using the BUSCO land plants (embryophyta) benchmark gene set (Supplementary Table 8). Whole-genome synteny comparison between Bartlett.DH_v2, the chromosome assembly of the Bartlett cultivar, and d’Anjou meta-assembly were evaluated with D-GENIES^98^ using repeat masked (http://www.repeatmasker.org) DNA alignments performed with minimap2^99^ for the whole genome and each of the 17 *Pyrus communis* chromosomes as shown in Fig. 1 and Supplementary Fig. 6, respectively.

### Gene prediction

To identify the regions of genomic DNA that encode genes, we first estimated the portion of d’Anjou meta-assembly comprised of repetitive elements suitable for repeat masking prior to protein-coding gene annotation following the protocol described by Campbell *et al* (2014)^100^. The meta-assembly was first searched using MITE-Hunter^101^ and LTRharvest/ LTRdigest^102, 103^ to collect consensus miniature inverted-repeat transposable elements (MITEs) and long terminal repeat retrotransposons (LTRs) respectively. LTRs were filtered to remove false positives and elements with nested insertions and used together with the MITEs to mask the genomes. The unmasked regions of the genomes were then annotated with RepeatModeler (http://www.repeatmasker.org/RepeatModeler) to predict additional *de novo* repetitive sequences. All collected repetitive sequences were compared to a BLAST database of plant proteins from SwissProt and RefSeq, and sequences with significant hits were excluded from the repeat masking library.

Extensive extrinsic gene annotation homology evidence from RNA-seq and protein were collected to supplement *ab initio* gene predictions. RNA-Seq evidence included Trinity^104^ *de novo* reconstructed transcripts from d’Anjou pear fruit peel and cortex tissues sampled at multiple time points described in our previous study^32^. Protein homology evidence of closely related species were collected from the Genome Database for Rosaceae (GDR), including *Malus domestica*, *Prunus persica*, *Pyrus betulifolia*, *Pyrus communis* ‘Bartlett’, *Pyrus* x *bretschneideri*, *Rosa chinensis*, and *Rubus occidentalis*^105^. The plant model species, *Arabidopsis thaliana*^106^, was included as well.

Protein-coding gene annotations from the *Pyrus communis* reference genomes of Bartlett_v1 and Bartlett.DH_v2 were separately transferred (liftovers) to pseudomolecules of d’Anjou meta-assembly using the FLO (https://github.com/wurmlab/flo) pipeline based on the UCSC Genome Browser Kent-Toolkit^107^. Next, the MAKER annotation pipeline (release 3.01.02)^108^ was used to update the transferred annotations with evidence data and gene models predicted by *ab initio* gene finders. Repetitive and low complexity regions of the pseudomolecules were first masked with RepeatMasker in MAKER using the previously described d’Anjou-specific repeat library. MAKER updated transferred annotations with evidence data and predicted additional annotations with Augustus^109, 110^ and SNAP^111^ using the d’Anjou training set where evidence suggests a gene. Only predicted gene models supported by annotation evidence, encode a Pfam domain, or both, were retained.

### Computation of pear orthogroups

To compare the gene content of the two *Pyrus communis* cultivars, ‘Bartlett’ and ‘d’Anjou’, orthologous and paralogous gene clusters of Bartlett_v1, Bartlett.DH_v2, and d’Anjou were estimated with OrthoFinder version 1.1.5^112^ for annotated proteins in all the genomes.

### Bartlett.DH_v2 genome polishing

To improve the base quality of the publicly available pear reference genome, the *Pyrus communis* ‘Bartlett.DH_v2’ assembly was iteratively polished with two rounds of Pilon (v1.24)^95^ using the raw Illumina shotgun reads from the Bartlett.DH_v2 genome projects obtained from the NCBI Short Read Archive (SRA accessions: SRR10030340, SRR10030308), and completeness and accuracy assessed with the BUSCO^30^ embryophyta_odb10 database.

### Gene family identification

Coding sequences of candidate genes and their corresponding peptides gleaned from published literature were sorted into pre-computed orthologous gene family clusters of representative 26 genomes from land plants using the both BLASTp^113^ and HMMER hmmscan^114^ sequence search option of the *GeneFamilyClassifier* tool implemented in the PlantTribes gene family analysis pipeline (https://github.com/dePamphilis/PlantTribes). Classification results, including orthogroup taxa gene counts, corresponding superclusters (super orthogroups) at multiple clustering stringencies, and orthogroup-level annotations from multiple public biological functional databases are reported in Supplementary Table 5.

### Gene family analysis

All the tools used in this process are modules from the command line version of PlantTribes software and are processed on SCINet (https://scinet.usda.gov/) with customized scripts. Protein coding genes from 14 Rosaceae species (*Fragaria vesca* v2.0.a2^115^, *Rosa chinensis* old Blush homozygous v2.0^116^, *Rubus occidentalis* v3.0^117^, *Prunus avium* v1.0.a1^118^, *Malus domestica* HFTH v1.0^13^, *M. domestica* GDDH13 v1.1^15^, *M. domestica* Gala v1.0^14^, *M. sieversii* v1.0^14^, *M. sylvestris* v1.0^14^, *Pyrus communis* v1.0^18^, *Pyrus communis* Bartlett DH v2.0^17^, *Pyrus ussuriensis* x *communis* v1.0^119^, *Pyrus bretschneideri* v1.1^120^, *Pyrus communis* d’Anjou v0.1) were sorted into orthologous groups with the *GeneFamilyClassifier* as previously described. For species lacking matching coding sequence file and peptide file, transcripts were processed to predict potential protein coding regions using the TransDecoder^121^ option of *AssemblyPostProcessor*. A detailed summary of the Rosaceae gene family classification results are in Supplementary Table 3. Sequences classified into the orthogroups of interest (with candidate genes in this study) were integrated with scaffold backbone gene models using the *GeneFamilyIntegrator* tool. Gene names were modified as shown in Supplementary Table 9 for easier recognition of the species. Amino acid multiple sequence alignments and their corresponding DNA codon alignments were generated by *GeneFamilyAligner* with the L-INS-i algorithm implemented in MAFFT^122^. Sites present in less than 10% of the aligned DNA sequences were removed with trimAL^123^. Maximum likelihood (ML) phylogenetic trees were estimated from the trimmed DNA alignments using the RAxML algorithm^124^ option in the *GeneFamilyPhylogenyBuilder*. One hundred bootstrap replicates (unless otherwise indicated) were conducted for each tree to estimate the reliability of the branches. The multiple sequence alignments were visualized in the Geneious R9 software^125^ with Clustal color scheme. The phylogeny was colored with a custom script and visualized with Dendroscope (version 3.7.5)^126^. Gene sequences, alignments, and phylogenies are available in Supplementary File 1-3.

### Domain prediction

To estimate domain structures of proteins in each orthogroup, the predicted amino acid sequences (either obtained from public databases or generated by the PlantTribes *AssemblyPostProcessor* tool) were submitted to interproscan (version 5.44-79.0)^127, 128^ on SCINet and searched against all the databases.

### Targeted gene family annotation

The following approaches were used in parallel to annotate candidate genes from the original Bartlett.DH_v2, polished Bartlett.DH_v2, and the d’Anjou genome assemblies:

#### TGFam-finder^67^

The ‘RESOURCE.config’ and ‘PROGRAM_PATH.config’ files were generated according to the author’s instruction. The two Bartlett.DH_v2 genome assemblies and the d’Anjou_v1 genome were used as the *target genomes*. Complete protein sequences from apples and pears in the same orthogroup were used as *protein for domain identification*. Complete protein sequences from other Rosaceae species and *Arabidopsis thaliana* in the same orthogroup were used as *resource proteins* for each annotation step. For each orthogroup, Pfam annotations from the InterProScan results were used as *TSV for domain identification*. For orthogroups without Pfam descriptions, MobiDBLite information was used as *TSV for domain identification*.

#### Bitacora^66^

Arabidopsis genes from targeted gene families (orthogroups of interest) were used to generate a multiple sequence alignment and HMM profile using MAFFT^122^ and hmmbuild. The resulting files were then used as input for Bitacora v1.3, running in both genome mode and full mode to identify genes of interest in the original Bartlett.DH_v2 genome.

### Manual curation and gene model verification

In cases where both TGFam-Finder and Bitacora failed to predict a full-length gene, the gene model was curated manually.

#### Curation with orthologous gene models

First, the genomic region containing the target sequence was determined either by the general feature format file (gff) or a BLASTn search using the coding sequence of the target gene or a closely related gene as a query. Next, a genomic fragment containing the target sequence and 3kb upstream and downstream of the targeted region was extracted. Then, the incomplete transcript(s), predicted exons, and complete gene models from a closely related species were mapped to the extracted genomic region. The final gene model was determined by using the full-length coding sequence of a closely related gene as a reference.

#### Curation with RNA-seq read mapping

The gff3 files obtained from Bitacora were loaded into an Apollo docker container (v2.6.3)^129^ for verification of the predicted gene models using expression data. Publicly available RNA-seq data^68–73^ for *Pyrus* were used as inputs of an RNA-seq aligner, STAR (v2.7.8a)^130^, and alignments were performed with maximum intron size set to 5kb and default settings. Intron-exon structure was compared to the aligned expression data. If there was insufficient RNA-seq coverage from the targeted cultivar, data from other cultivars and *Pyrus* species were used as supporting evidence. Read mapping results are available in Supplementary File 4-5. Curated gene models from the original Bartlett.DH_v2 were transferred to the polished genome for validation.

Gene model cartoons were generated using the visualize gene structure function in TBtools (v1.09854)^131^. Final gene models and their corresponding chromosomal locations are available in Supplementary File 6-7.

### Data Availability

Raw read data of d’Anjou genome sequencing has been deposited at NCBI SRA under Bioproject ID PRJNA762155. Genome assembly and gene prediction of the draft ‘d’Anjou’ genome, and the polished ‘Bartlett.DH_v2’ genome assembly have been submitted to the Genome Database for Rosacea (GDR). Supplementary information accompanies the manuscript on the BioRxiv.

## Supporting information

README

## Acknowledgements

This work was supported by the Washington Tree Fruit Research Commission project PR-17-104 and the Agricultural Research Service in the US Department of Agriculture. The authors would like to acknowledge Heidi Hargarten for maintaining the d’Anjou plant and collecting leaf tissue for sequencing. They also thank Craig Praul at Penn State and Diana Burkart-Waco and Lutz Froenicke at UC Davis for sequencing.

## Author Contributions

HZ, JW, LH conceived and designed the research. PR prepared gDNA for sequencing. HZ, EW, PT, JE, JW, and AH performed the genome assembly and gene family analysis. All authors participated in writing and revising the manuscript.

## References

1. Chen, J. et al. A chromosome-scale genome sequence of pitaya (*Hylocereus undatus*) provides novel insights into the genome evolution and regulation of betalain biosynthesis. Hortic Res. 8, 164 (2021).

2. Wang, J. et al. A high-quality genome assembly of *Morinda officinalis*, a famous native southern herb in the Lingnan region of southern China. Hortic Res. 8, 135 (2021).

3. Huang, Y. et al. Genome of a citrus rootstock and global DNA demethylation caused by heterografting. Hortic Res. 8, 69 (2021).

4. Chen, D. et al. The chromosome-level reference genome of *Coptis chinensis* provides insights into genomic evolution and berberine biosynthesis. Hortic Res. 8: 121 (2021).

5. Xu, X. et al. The chromosome-level *Stevia* genome provides insights into steviol glycoside biosynthesis. Hortic Res. 8: 129 (2021).

6. Wang, P. et al. Genetic basis of high aroma and stress tolerance in the oolong tea cultivar genome. Hortic Res. 8: 107 (2021).

7. Hill, J. L. & Hollender, C. A. Branching out: new insights into the genetic regulation of shoot architecture in trees. Curr Opin Plant Biol. 47: 73–80 (2019).

8. Stansell, Z. & Björkman, T. From landrace to modern hybrid broccoli: the genomic and morphological domestication syndrome within a diverse *B. oleracea* collection. Hortic Res. 7: 159 (2020).

9. Stansell, Z. et al. Genotyping-by-sequencing of *Brassica oleracea* vegetables reveals unique phylogenetic patterns, population structure and domestication footprints. Hortic Res. 5: 38 (2018).

10. Cheng, F. et al. Subgenome parallel selection is associated with morphotype diversification and convergent crop domestication in *Brassica rapa* and *Brassica oleracea*. Nat Genet. 48: 1218–1224 (2016).

11. Mabry, M. E. et al. The evolutionary history of wild, domesticated, and feral *Brassica Oleracea* (Brassicaceae). Mol Biol Evol. doi:10.1093/molbev/msab183 (2021).

12. Cheng, F. et al. Genome resequencing and comparative variome analysis in a *Brassica rapa* and *Brassica oleracea* collection. Sci Data. 3: 160119 (2016).

13. Zhang, L. et al. A high-quality apple genome assembly reveals the association of a retrotransposon and red fruit colour. Nat Commun. 10: 1494 (2019).

14. Sun, X. et al. Phased diploid genome assemblies and pan-genomes provide insights into the genetic history of apple domestication. Nat Genet. 52: 1423–1432 (2020).

15. Daccord, N. et al. High-quality *de novo* assembly of the apple genome and methylome dynamics of early fruit development. Nat Genet. 49: 1099–1106 (2017).

16. Velasco, R. et al. The genome of the domesticated apple (*Malus × domestica* Borkh.). Nat Genet. 42: 833–839 (2010).

17. Linsmith, G. et al. Pseudo-chromosome-length genome assembly of a double haploid “Bartlett” pear (*Pyrus communis* L.). Gigascience. 8. doi:10.1093/gigascience/giz138 (2019).

18. Chagné, D. et al. The draft genome sequence of European pear (*Pyrus communis* L. ‘Bartlett’). Plos One. 9: e92644 (2014).

19. Wu, J. et al. A naturally occurring InDel variation in *BraA*.FLC.b (BrFL*C2*) associated with flowering time variation in *Brassica rapa*. Bmc Plant Biol. 12: 151 (2012).

20. Tollenaere, R. et al. Identification and characterization of candidate *Rlm4* blackleg resistance genes in *Brassica napus* using nextLJgeneration sequencing. Plant Biotechnol J. 10: 709–715 (2012).

21. Takos, A. M. et al. Light-Induced Expression of a *MYB* Gene Regulates Anthocyanin Biosynthesis in Red Apples. Plant Physiology 142: 1216–1232 (2006).

22. Pertea, M. et al. CHESS: a new human gene catalog curated from thousands of large-scale RNA sequencing experiments reveals extensive transcriptional noise. Genome Biol 19: 208 (2018).

23. Pilkington, S. M. et al. A manually annotated *Actinidia chinensis* var. *chinensis* (kiwifruit) genome highlights the challenges associated with draft genomes and gene prediction in plants. Bmc Genomics 19: 257 (2018).

24. Marx, H. et al. A proteomic atlas of the legume *Medicago truncatula* and its nitrogen-fixing endosymbiont *Sinorhizobium meliloti*. Nat Biotechnol. 34: 1198–1205 (2016).

25. Kyriakidou, M., Tai, H. H., Anglin, N. L., Ellis, D., & Strömvik, M. V. Current strategies of polyploid plant genome sequence assembly. Front Plant Sci. 9: 1660 (2018).

26. Hughes, T. E., Langdale, J. A., & Kelly, S. The impact of widespread regulatory neofunctionalization on homeolog gene evolution following whole-genome duplication in maize. Genome Res. 24: 1348–1355 (2014).

27. Yoshida, S. et al. Genome sequence of *Striga asiatica* provides insight into the evolution of plant parasitism. Curr Biol. 29: 3041–3052.e4 (2019).

28. Yang, Z. et al. Comparative transcriptome analyses reveal core parasitism genes and suggest gene duplication and repurposing as sources of structural novelty. Mol Biol Evol. 32: 767–790 (2015).

29. Xiao, D. et al. The *Brassica rapa FLC* homologue *FLC2* is a key regulator of flowering time, identified through transcriptional co-expression networks. J Exp Bot. 64: 4503–4516 (2013).

30. Manni, M., Berkeley, M. R., Seppey, M., Simão, F. A., & Zdobnov, E. M. BUSCO update: novel and streamlined workflows along with broader and deeper phylogenetic coverage for scoring of eukaryotic, prokaryotic, and viral genomes. Mol Biol Evol. msab199-(2021).

31. Mistry, J. et al. Pfam: The protein families database in 2021. Nucleic Acids Res. 49: gkaa913- (2020).

32. Honaas, L. et al. Transcriptomics of differential ripening in ‘d’Anjou’ pear (*Pyrus communis* L.). Front Plant Sci. 12: 609684 (2021).

33. Kwon, M. & Choe, S. Brassinosteroid biosynthesis and dwarf mutants. J Plant Biol. 48: 1 (2004).

34. Waite, J. M. & Dardick, C. The roles of the *IGT* gene family in plant architecture: past, present, and future. Curr Opin Plant Biol. 59: 101983 (2021).

35. Bulley, S. M. et al. Modification of gibberellin biosynthesis in the grafted apple scion allows control of tree height independent of the rootstock. Plant Biotechnol J. 3: 215–223 (2005).

36. Griffiths, J. et al. Genetic Characterization and Functional Analysis of the *GID1* Gibberellin Receptors in Arabidopsis. Plant Cell Online 18: 3399–3414 (2006).

37. Dugardeyn, J. & Straeten, D. V. D. Ethylene: Fine-tuning plant growth and development by stimulation and inhibition of elongation. Plant Sci. 175: 59–70 (2008).

38. K e ek, P. et al. The PIN-FORMED (PIN) protein family of auxin transporters. Genome ř č Biol. 10: 249 (2009).

39. Martín Rejano, E. M. et al. Auxin and ethylene are involved in the responses of root system LJ architecture to low boron supply in Arabidopsis seedlings. Physiol Plantarum. 142: 170–178 (2011).

40. Li, H. L. et al. Possible roles of auxin and zeatin for initiating the dwarfing effect of M9 used as apple rootstock or interstock. Acta Physiol Plant. 34: 235–244 (2012).

41. Chang, L., Ramireddy, E., & Schmülling, T. Lateral root formation and growth of Arabidopsis is redundantly regulated by cytokinin metabolism and signalling genes. J Exp Bot. 64: 5021–5032 (2013).

42. Dardick, C. et al. *PpeTAC1* promotes the horizontal growth of branches in peach trees and is a member of a functionally conserved gene family found in diverse plants species. Plant J. 75: 618–630 (2013).

43. Tworkoski, T., Webb, K., & Callahan, A. Auxin levels and *MAX1–4* and *TAC1* gene expression in different growth habits of peach. Plant Growth Regul. 77: 279–288 (2015).

44. Hollender, C. A., Hadiarto, T., Srinivasan, C., Scorza, R., & Dardick, C. A brachytic dwarfism trait (dw) in peach trees is caused by a nonsense mutation within the gibberellic acid receptor PpeGID1c. New Phytol. 210: 227–239 (2016).

45. Ma, Y. et al. Involvement of auxin and brassinosteroid in dwarfism of autotetraploid apple (*Malus × domestica*). Sci Rep-uk. 6: 26719 (2016).

46. Li, G. et al. Transcriptome analysis reveals the effects of sugar metabolism and auxin and cytokinin signaling pathways on root growth and development of grafted apple. Bmc Genomics. 17: 150 (2016).

47. Foster, T. M., McAtee, P. A., Waite, C. N., Boldingh, H. L., & McGhie, T. K. Apple dwarfing rootstocks exhibit an imbalance in carbohydrate allocation and reduced cell growth and metabolism. Hortic Res. 4: 17009 (2017).

48. Harrison C. J. Auxin transport in the evolution of branching forms. New Phytol. 215: 545– 551 (2017).

49. Guseman, J. M., Webb, K., Srinivasan, C., & Dardick, C. *DRO1* influences root system architecture in Arabidopsis and *Prunus* species. Plant J. 89: 1093–1105 (2017).

50. An, H. et al. Dwarfing effect of apple rootstocks is intimately associated with low number of fine roots. Hortscience. 52: 503–512 (2017).

51. An, J. et al. Molecular cloning and functional characterization of *MdPIN1* in apple. J Integr Agr. 16: 1103–1111 (2017).

52. Zheng, L. et al. Genome-wide identification and expression analysis of brassinosteroid biosynthesis and metabolism genes regulating apple tree shoot and lateral root growth. J Plant Physiol. 231: 68–85 (2018).

53. Gan, Z. et al. *MdPIN1b* encodes a putative auxin efflux carrier and has different expression patterns in BC and M9 apple rootstocks. Plant Mol Biol. 96: 353–365 (2018).

54. Zheng, X. et al. *MdWRKY9* overexpression confers intensive dwarfing in the M26 rootstock of apple by directly inhibiting brassinosteroid synthetase *MdDWF4* expression. New Phytol. 217: 1086–1098 (2018).

55. Wang, L. et al. The isolation of the *IGT* family genes in *Malus × domestica* and their expressions in four idiotype apple cultivars. Tree Genet Genomes. 14: 46 (2018).

56. Cheng, J. et al. A single nucleotide mutation in *GID1c* disrupts its interaction with *DELLA1* and causes a GA insensitive dwarf phenotype in peach. Plant Biotechnol J. 17: 1723–1735 LJ (2019).

57. Li, C. et al. Comprehensive expression analysis of Arabidopsis *GA2-oxidase* genes and their functional insights. Plant Sci. 285: 1–13 (2019).

58. Zheng, X., Zhang, H., Xiao, Y., Wang, C., & Tian, Y. Deletion in the promoter of *PcPIN-L* affects the polar auxin transport in dwarf pear (*Pyrus communis* L.). Sci Rep-uk. 9: 18645 (2019).

59. Swarup, R. & Bhosale, R. Developmental roles of AUX1/LAX auxin influx carriers in plants. Front Plant Sci. 10: 1306 (2019).

60. Tan, M. et al. Role of cytokinin, strigolactone, and auxin export on outgrowth of axillary buds in apple. Front Plant Sci. 10: 616 (2019).

61. Qi, L. et al. Characterization of the auxin efflux transporter PIN proteins in pear. Plants. 9: 349 (2020).

62. Sun, X. et al. Genes involved in strigolactone biosyntheses and their expression analyses in columnar apple and standard apple. Biol Plantarum. 64: 68–76 (2020).

63. Cheng, J. et al. Functional analysis of the *Gibberellin 2-oxidase* gene family in peach. Front Plant Sci. 12: 619158 (2021).

64. Pandey, B. K. et al. Plant roots sense soil compaction through restricted ethylene diffusion. Science 371: 276–280 (2021).

65. Wafula E. K. Computational methods for comparative genomics of non-model species: A case study in the parasitic plant family Orobanchaceae. (The Pennsylvania State University, State College, PA, USA, 2019).

66. Vizueta, J., Sánchez Gracia, A., & Rozas, J. bitacora: A comprehensive tool for the LJ identification and annotation of gene families in genome assemblies. Mol Ecol Resour. 20: 1445–1452 (2020).

67. Kim, S. et al. TGFam Finder: a novel solution for target gene family annotation in plants. LJ LJ New Phytol. 227: 1568–1581 (2020).

68. Nham, N. T. et al. A transcriptome approach towards understanding the development of ripening capacity in ‘Bartlett’ pears (*Pyrus communis* L.). Bmc Genomics. 16: 762 (2015).

69. Hewitt, S. L., Hendrickson, C. A., & Dhingra, A. Evidence for the involvement of vernalization-related genes in the regulation of cold-induced ripening in ‘d’Anjou’ and ‘Bartlett’ pear fruit. Sci Rep-uk. 10: 8478 (2020).

70. Gabay, G. et al. Transcriptome analysis and metabolic profiling reveal the key role of -α linolenic acid in dormancy regulation of European pear. J Exp Bot. 70: ery405 (2018).

71. Zhang, Z. et al. Transcriptomic and metabolomic analysis provides insights into anthocyanin and procyanidin accumulation in pear. Bmc Plant Biol. 20: 129 (2020).

72. Nham, N. T., Macnish, A. J., Zakharov, F., & Mitcham, E. J. ‘Bartlett’ pear fruit (*Pyrus communis* L.) ripening regulation by low temperatures involves genes associated with jasmonic acid, cold response, and transcription factors. Plant Sci. 260: 8–18 (2017).

73. Zhang, H. et al. RNA-Seq analysis of the tissue-specific expressed genes of *Pyrus betulaefolia* in root, stem and leaf. Acta Horticulturae Sinica. 45: 1881–1894 (2018).

74. Yoshihara, T., Spalding, E. P., & Iino, M. *AtLAZY1* is a signaling component required for gravitropism of the *Arabidopsis thaliana* inflorescence. Plant J. 74: 267–279 (2013).

75. Yoshihara, T., & Spalding, E. P. *LAZY* genes mediate the effects of gravity on auxin gradients and plant architecture. Plant Physiol. 175: 959–969 (2017).

76. Li, P. et al. *LAZY1* controls rice shoot gravitropism through regulating polar auxin transport. Cell Res. 17: 402–410 (2007).

77. Yu, B. et al. *TAC1*, a major quantitative trait locus controlling tiller angle in rice. Plant J. 52: 891–898 (2007).

78. Uga, Y. et al. Control of root system architecture by *DEEPER ROOTING 1* increases rice yield under drought conditions. Nat Genet. 45: 1097–1102 (2013).

79. Mount, S. M. Genomic sequence, splicing, and gene annotation. Am J Hum Genetics. 67: 788–792 (2000).

80. Guo, L., & Liu, C-M. A single-nucleotide exon found in Arabidopsis. Sci Rep-uk. 5: 18087 (2015).

81. Sharma, S., Sharma, S. N., & Saxena, R. Identification of short exons disunited by a short intron in eukaryotic DNA regions. Ieee Acm Transactions Comput Biology Bioinform. 17: 1660–1670 (2018).

82. Zheng, X., Xiao, Y., Tian, Y., Yang, S., & Wang, C. *PcDWF1*, a pear brassinosteroid biosynthetic gene homologous to *AtDWARF1*, affected the vegetative and reproductive growth of plants. Bmc Plant Biol. 20: 109 (2020).

83. Cancino-García, V. J., Ramírez-Prado, J. H., & De-la-Peña, C. Auxin perception in *Agave* is dependent on the species’ Auxin Response Factors. Sci Rep-uk. 10: 3860 (2020).

84. Feng, Y. et al. Genome-wide identification and characterization of ABC transporters in nine Rosaceae species identifying *MdABCG28* as a possible cytokinin transporter linked to dwarfing. Int J Mol Sci. 20: 5783 (2019).

85. Wall, P. K. et al. PlantTribes: a gene and gene family resource for comparative genomics in plants. Nucleic Acids Res. 36: D970–D976 (2008).

86. Harper, L. et al. AgBioData consortium recommendations for sustainable genomics and genetics databases for agriculture. Database. 2018: 1–32 (2018).

87. Michiels, A., Ende, W. V. den., Tucker, M., Riet, L. V., & Laere, A. V. Extraction of high-quality genomic DNA from latex-containing plants. Anal Biochem. 315: 85–89 (2003).

88. Koren, S. et al. Canu: scalable and accurate long-read assembly via adaptive k-mer weighting and repeat separation. Genome Res. 27: 722–736 (2017).

89. Zimin, A. V. et al. The MaSuRCA genome assembler. Bioinformatics. 29: 2669–2677 (2013).

90. Weisenfeld, N. I., Kumar, V., Shah, P., Church, D. M., & Jaffe, D. B. Direct determination of diploid genome sequences. Genome Res. 27: 757–767 (2017).

91. Chakraborty, M., Baldwin-Brown, J. G., Long, A. D., & Emerson, J. J. Contiguous and accurate *de novo* assembly of metazoan genomes with modest long read coverage. Nucleic Acids Res. 44: e147–e147 (2016).

92. Jackman, S. D. et al. Tigmint: correcting assembly errors using linked reads from large molecules. Bmc Bioinformatics. 19: 393 (2018).

93. Yeo, S., Coombe, L., Warren, R. L., Chu, J., & Birol, I. ARCS: scaffolding genome drafts with linked reads. Bioinformatics. 34: 725–731 (2017).

94. Boetzer, M. & Pirovano, W. Toward almost closed genomes with GapFiller. Genome Biol. 13: R56 (2012).

95. Walker, B. J. et al. Pilon: An integrated tool for comprehensive microbial variant detection and genome assembly improvement. Plos One. 9: e112963 (2014).

96. Hämälä, T. et al. Genomic structural variants constrain and facilitate adaptation in natural populations of *Theobroma cacao*, the chocolate tree. PNAS. doi:10.1073/pnas.2102914118 (2021).

97. Alonge, M. et al. RaGOO: fast and accurate reference-guided scaffolding of draft genomes. Genome Biol. 20: 224 (2019).

98. Cabanettes, F. & Klopp, C. D-GENIES: dot plot large genomes in an interactive, efficient and simple way. Peerj. 6: e4958 (2018).

99. Li H. Minimap and miniasm: fast mapping and *de novo* assembly for noisy long sequences. Bioinformatics. 32: 2103–2110 (2016).

100. Campbell, M. S., Holt, C., Moore, B., & Yandell, M. Genome annotation and curation using MAKER and MAKERLJP. Curr Protoc Bioinform. 48: 4.11.1-4.11.39 (2014).

101. Han, Y. & Wessler, S. R. MITE-Hunter: a program for discovering miniature inverted-repeat transposable elements from genomic sequences. Nucleic Acids Res. 38: e199–e199 (2010).

102. Ellinghaus, D., Kurtz, S., & Willhoeft, U. LTRharvest, an efficient and flexible software for de novo detection of LTR retrotransposons. Bmc Bioinformatics. 9: 18 (2008).

103. Steinbiss, S., Willhoeft, U., Gremme, G., & Kurtz, S. Fine-grained annotation and classification of *de novo* predicted LTR retrotransposons. Nucleic Acids Res. 37: 7002–7013 (2009).

104. Grabherr, M. G. et al. Full-length transcriptome assembly from RNA-Seq data without a reference genome. Nat Biotechnol. 29: 644–652 (2011).

105. Jung, S. et al. 15 years of GDR: New data and functionality in the Genome Database for Rosaceae. Nucleic Acids Res. 47: D1137–D1145 (2018).

106. Cheng, C. et al. Araport11: a complete reannotation of the *Arabidopsis thaliana* reference genome. Plant J. 89: 789–804 (2017).

107. Kuhn, R. M., Haussler, D., & Kent, W. J. The UCSC genome browser and associated tools. Brief Bioinform. 14: 144–161 (2013).

108. Cantarel, B. L. et al. MAKER: An easy-to-use annotation pipeline designed for emerging model organism genomes. Genome Res. 18: 188–196 (2008).

109. Stanke, M., Steinkamp, R., Waack, S., & Morgenstern, B. AUGUSTUS: a web server for gene finding in eukaryotes. Nucleic Acids Res. 32: W309–W312 (2004).

110. Hoff, K. J. & Stanke, M. Predicting genes in single genomes with AUGUSTUS. Curr Protoc Bioinform. 65: e57 (2019).

111. Korf, I. Gene finding in novel genomes. Bmc Bioinformatics. 5: 59 (2004).

112. Emms, D. M. & Kelly, S. OrthoFinder: phylogenetic orthology inference for comparative genomics. Genome Biol. 20: 238 (2019).

113. Camacho, C. et al. BLAST+: architecture and applications. Bmc Bioinformatics. 10: 421 (2009).

114. Eddy, S. R. Accelerated Profile HMM Searches. Plos Comput Biol. 7: e1002195 (2011).

115. Li, Y. et al. Genome re-annotation of the wild strawberry Fragaria vesca using extensive Illumina- and SMRT-based RNA-seq datasets. Dna Res. 25: dsx038 (2017).

116. Raymond, O. et al. The *Rosa* genome provides new insights into the domestication of modern roses. Nat Genet. 50: 772–777 (2018).

117. VanBuren, R. et al. A near complete, chromosome-scale assembly of the black raspberry (*Rubus occidentalis*) genome. Gigascience. 7: giy094- (2018).

118. Shirasawa, K. et al. The genome sequence of sweet cherry (*Prunus avium*) for use in genomics-assisted breeding. Dna Res. 24: dsx020- (2017).

119. Ou, C. et al. A *de novo* genome assembly of the dwarfing pear rootstock Zhongai 1. Sci Data 6: 281 (2019).

120. Xue, H. et al. Chromosome level high-density integrated genetic maps improve the *Pyrus bretschneideri* ‘DangshanSuli’ v1.0 genome. Bmc Genomics. 19: 833 (2018).

121. Haas, B. J. et al. *De novo* transcript sequence reconstruction from RNA-seq using the Trinity platform for reference generation and analysis. Nat Protoc. 8: 1494–1512 (2013).

122. Katoh, K., Misawa, K., Kuma, K., & Miyata, T. MAFFT: a novel method for rapid multiple sequence alignment based on fast Fourier transform. Nucleic Acids Res. 30: 3059–3066 (2002).

123. Capella-Gutiérrez, S., Silla-Martínez, J. M., & Gabaldón, T. trimAl: a tool for automated alignment trimming in large-scale phylogenetic analyses. Bioinformatics. 25: 1972–1973 (2009).

124. Stamatakis, A. RAxML version 8: a tool for phylogenetic analysis and post-analysis of large phylogenies. Bioinformatics. 30: 1312–1313 (2014).

125. Kearse, M. et al. Geneious Basic: An integrated and extendable desktop software platform for the organization and analysis of sequence data. Bioinformatics. 28: 1647–1649 (2012).

126. Huson, D. H., & Scornavacca, C. Dendroscope 3: An interactive tool for rooted phylogenetic trees and networks. Systematic Biol. 61: 1061–1067 (2012).

127. Mulder, N. & Apweiler, R. Comparative Genomics. Methods Mol Biology. 396: 59–70 (2007).

128. Quevillon, E. et al. InterProScan: protein domains identifier. Nucleic Acids Res. 33: W116– W120 (2005).

129. Dunn, N. A. et al. Apollo: Democratizing genome annotation. Plos Comput Biol. 15: e1006790 (2019).

130. Dobin, A. et al. STAR: ultrafast universal RNA-seq aligner. Bioinformatics. 29: 15–21 (2013).

131. Chen, C. et al. TBtools: An integrative toolkit developed for interactive analyses of big biological data. Mol Plant. 13: 1194–1202 (2020).

