## Supplementary material for "Towards a catalog of pome tree architecture genes: the draft ‘d’Anjou’ genome (*Pyrus communis* L.)": README

README
Supplementary file 1: contains coding sequences and peptide sequences of all the members in each orthogroups in this study. Can be visualized in any text editor.
Supplementary file 2: contains the codon alignment and amino acid alignment of all the members in each orthogroups in this study. Can be visualized in any sequence alignment visulization tools.
Supplementary file 3: Newick format phylogenetic tree of all the orthogroups in this study. Can be visualized in any phylogenetic tree viewers.
Supplementary file 4: RNA-seq read mapping results. Six datasets from public databases mapped to the original Bartlett.DH_v2 assembly. Can be visualized in text editor.
Supplementary file 5: RNA-seq read mapping results. Six datasets from public databases mapped to the polished Bartlett.DH_v2 assembly. Can be visualized in text editor.
Supplementary file 6 and 7: curated gene models of Bartlett.DH_v2 and d'Anjou, respectively. Can be visualized in any text editor.
Supplenemtary file 8: scripts.
