## Supplemental figures for "Towards a catalog of pome tree architecture genes: the draft ‘d’Anjou’ genome (*Pyrus communis* L.)"

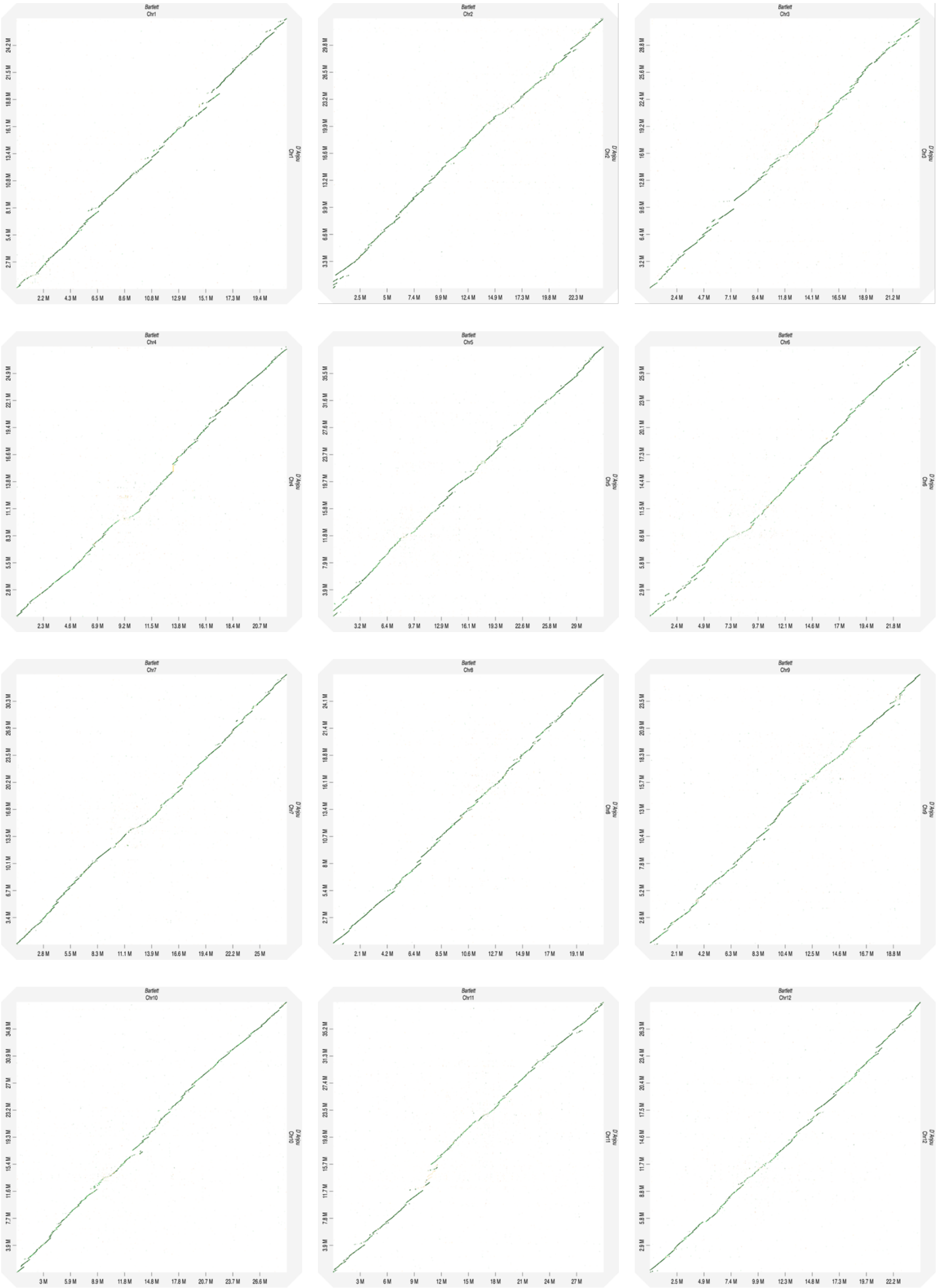
Bartlett
Chr1
D'Anjou
Chr1
Bartlett
Chr2
D'Anjou
Chr2
Bartlett
Chr3
D'Anjou
Chr3
Bartlett
Chr4
D'Anjou
Chr4
Bartlett
Chr5
D'Anjou
Chr5
Bartlett
Chr6
D'Anjou
Chr6
Bartlett
Chr7
D'Anjou
Chr7
Bartlett
Chr8
D'Anjou
Chr8
Bartlett
Chr9
D'Anjou
Chr9
Bartlett
Chr10
D'Anjou
Chr10
Bartlett
Chr11
D'Anjou
Chr11
Bartlett
Chr12
D'Anjou
Chr12

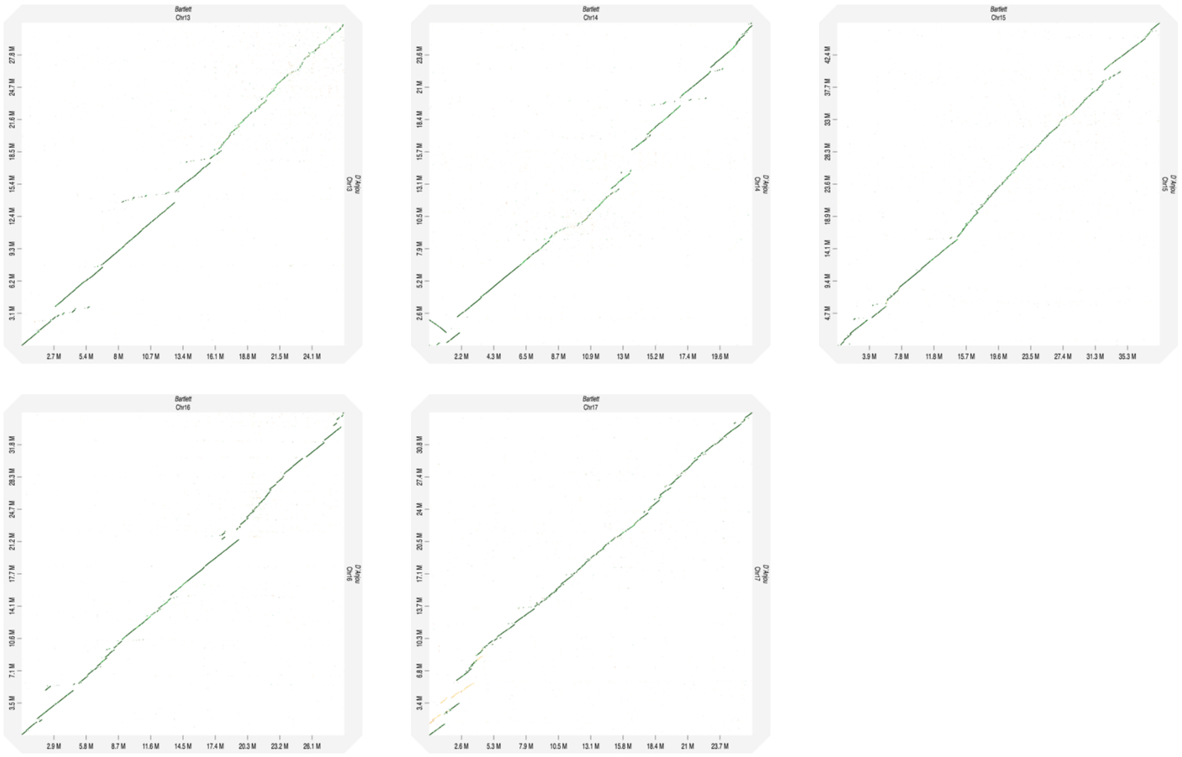

Supplementary Figure 1. Dot plot of chromosome alignments of Bartlett.DH_v2 (x axis) and d'Anjou_v1 (y axis).

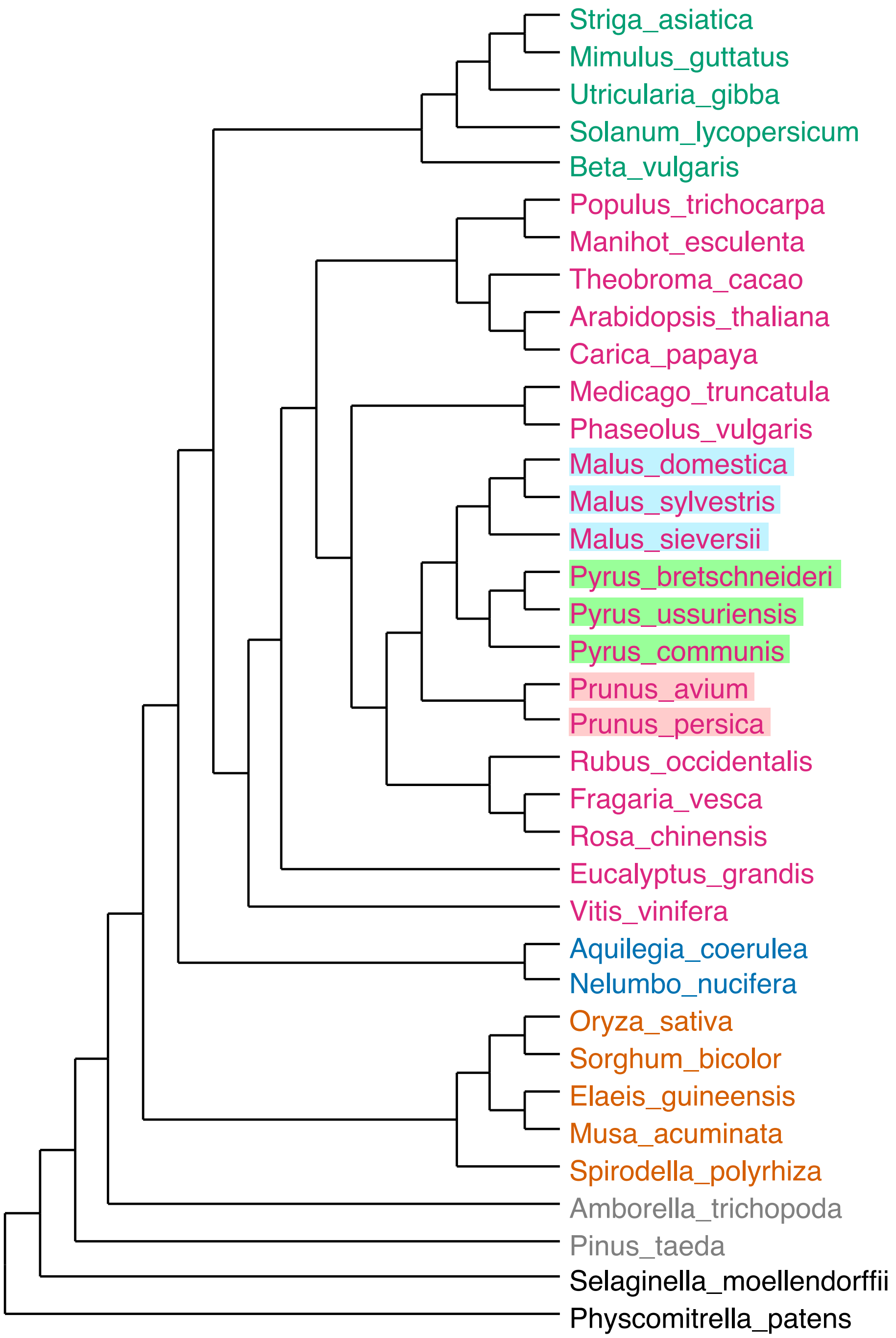


Supplementary Figure 2. A cladogram of the plant species used in this study. Rosaceae species except *Prunus persica* were classified into the 26 genome scaffold version 2. Genes from these species followed the same color scheme in the phylogenetic trees in this manuscript.

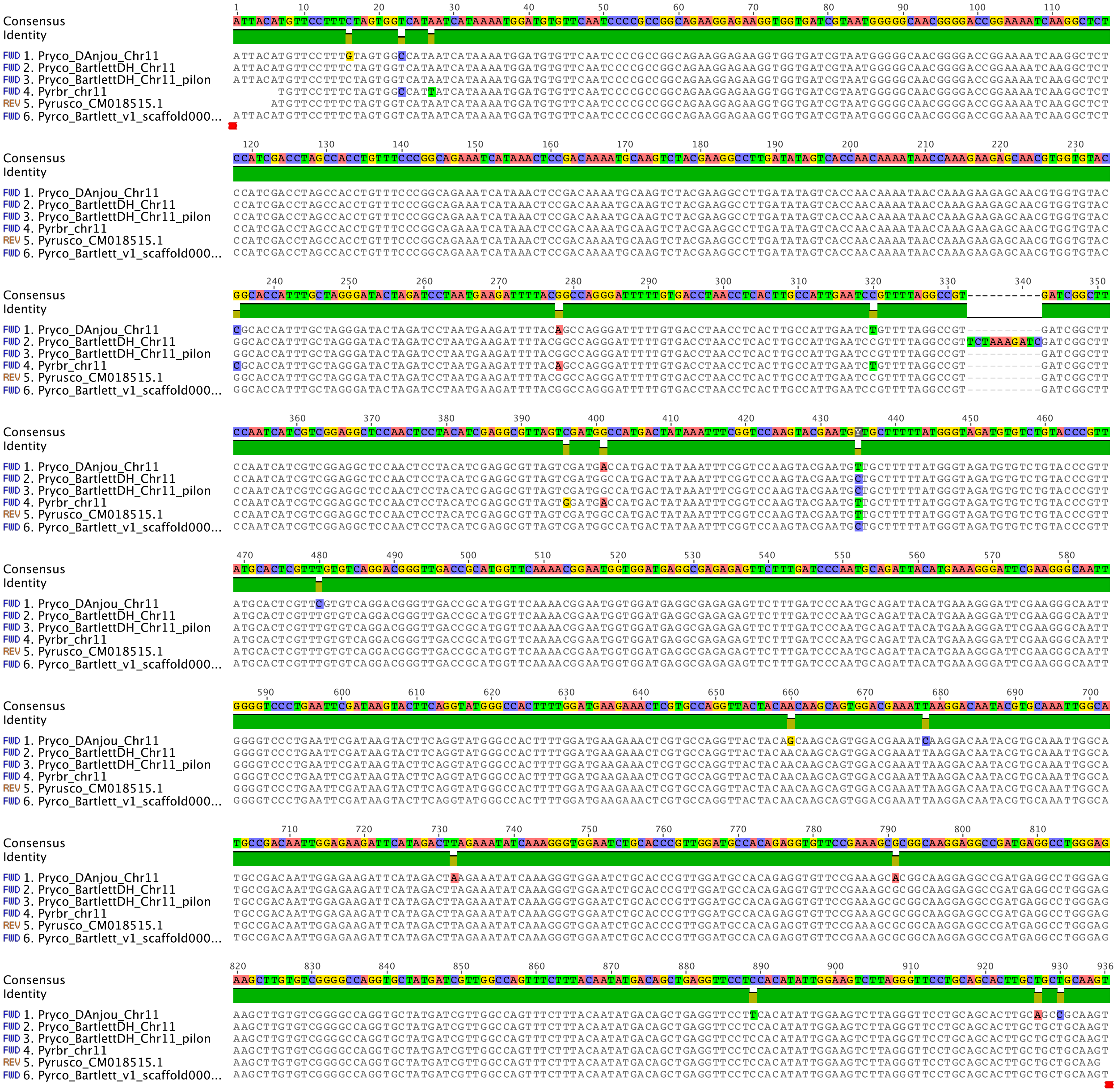


Supplementary Figure 3. Sequence comparison of genomic fragments where an *IPT* gene is expected to be located in various *Pyrus* genomes. Pyrco: *Pyrus communis*; Pyrbr: *P. bretschneideri*; Pyrusco: *P. ussuriensis x communis*. Sequence 2 is the original Bartlett.DH_v2 assembly and Sequence 3 is the polished assembly.

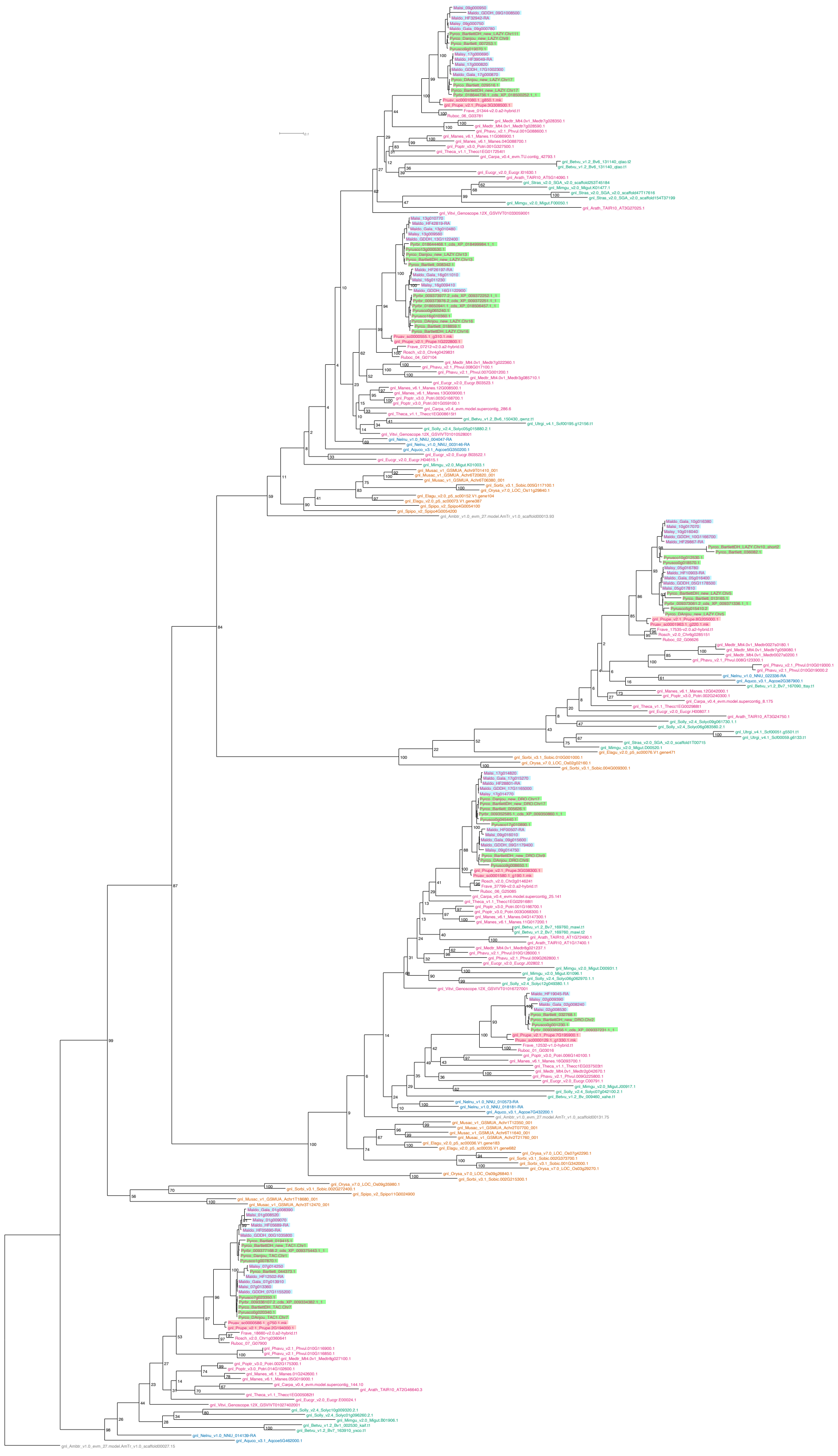

Supplementary Figure 4. Phylogram of the *IGT* gene family. Same tree as Fig 4a.

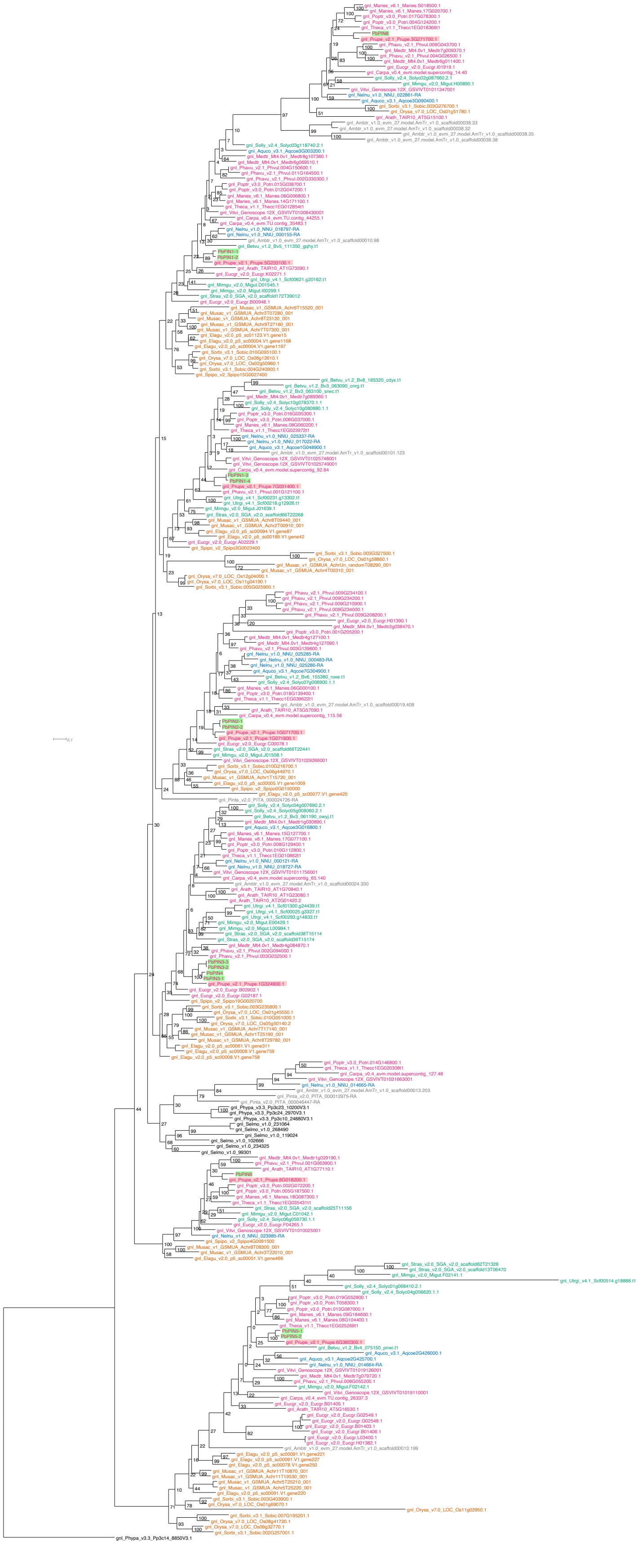

Supplementary Figure 5. Phylogeny of the *PIN* gene family inferred with the 26 genomes scaffold and *P. bretschneideri PIN* genes identified in reference 58.

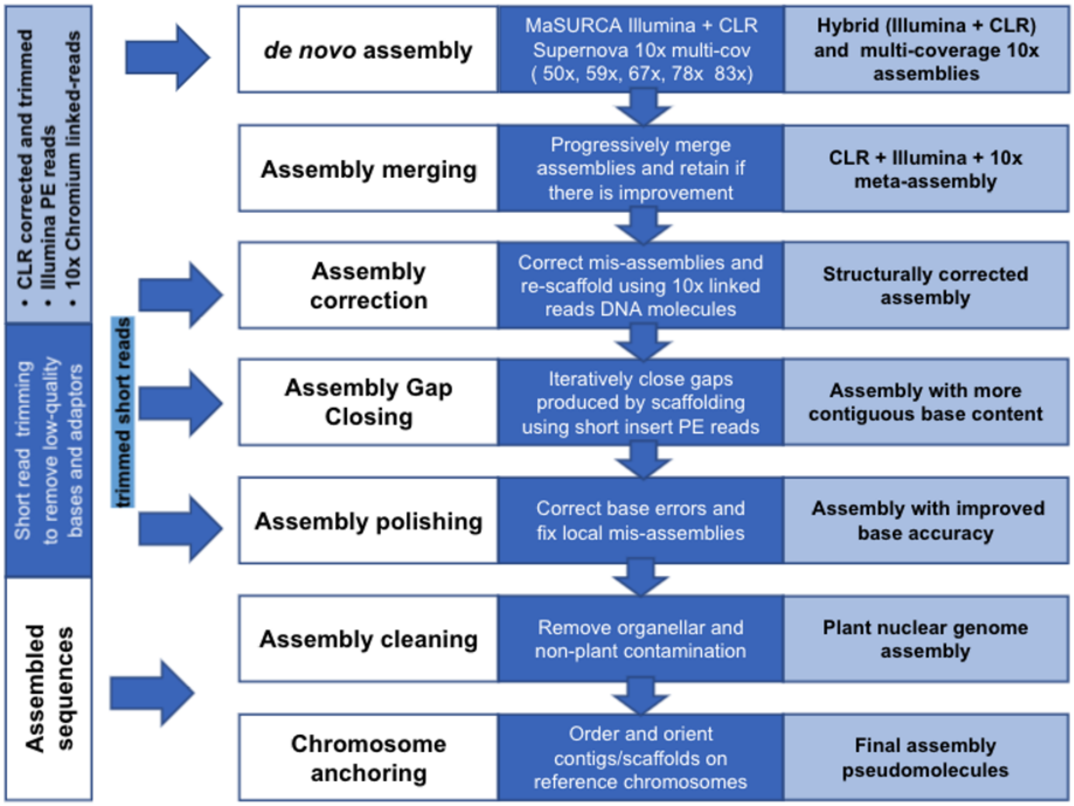


Supplementary Figure 6. A summary of the d’Anjou genome assembly pipeline.
